# Dynamic Coupling and Entropy Changes in KRAS G12D Mutation: Insights into Molecular Flexibility, Allostery and Function

**DOI:** 10.1101/2024.12.28.630590

**Authors:** Aysima Hacisuleyman, Deniz Yuret, Burak Erman

**Affiliations:** Department of Computational Biology, University of Lausanne, CH-1015 Lausanne, Switzerland; Department of Computer Engineering, Koc University Rumelifeneri Yolu, Sariyer 34450, Istanbul Turkey; Department of Chemical and Biological Engineering, Koc University Rumelifeneri Yolu, Sariyer 34450, Istanbul Turkey

**Keywords:** Mutual Information, Dynamic Allostery, Dynamic coupling, Kernel density estimation, Molecular dynamics

## Abstract

The oncogenic G12D mutation in KRAS is a major driver of cancer progression, yet the complete mechanism by which this mutation alters protein dynamics and function remains incompletely understood. Here, we investigate how the G12D mutation alters KRAS’s conformational landscape and residue-residue interactions using molecular dynamics simulations coupled with entropy calculations and mutual information (MI) analysis. We demonstrate that the mutation increases local entropy at key functional residues (D12, Y32, G60, and Q61), and introduces new peaks to the Ramachandran angles, disrupting the precise structural alignment necessary for GTP hydrolysis. Notably, while individual residue entropy increases, joint entropy analysis shows a complex reorganization pattern. MI analysis identifies enhanced dynamic coupling between distant residues, suggesting that the mutation establishes new long-range interactions that stabilize the active state. These findings show how G12D mutation redefines KRAS’s dynamic network, leading to persistent activation through enhanced residue coupling rather than mere local disruption. Our results suggest novel therapeutic strategies focused on modulating protein dynamics rather than targeting specific binding sites, potentially offering new approaches to combat KRAS-driven cancers.

**Highlights:**

- The G12D mutation increases entropy and mutual information between residue pairs, resulting in more correlated motions due to enhanced flexibility.
- Characteristic features of local and long-range allostery in wild-type (WT) and G12D KRAS are determined using structural proximity and mutual information analysis.
- The dynamics of KRAS interactions with its natural ligand atoms and the coordination of bridging water molecules provide insights into the changes induced by the G12D mutation.

## Introduction

KRAS, a pivotal regulator of cellular signaling pathways, is frequently mutated in various cancers, with the G12D mutation being an important example. This mutation significantly alters KRAS’s regulatory mechanisms, leading to the persistent activation of signaling pathways. One of KRAS’s critical functions is the hydrolysis of GTP to GDP, a process essential for turning off its signaling activity. However, the G12D mutation impairs this hydrolysis, resulting in continuous activation[1].

Mechanistic insights into KRAS function and dynamics have been significantly advanced through molecular dynamics (MD) simulations, which have clarified key structural features and conformational changes associated with its signaling mechanisms[2–7]. Molecular dynamics studies have shown that the malfunctioning of KRAS in the G12D mutant is driven by the increased flexibility and mobility of key residues due to the mutation[6, 8, 9]. The aspartic acid, Asp, group replacing glycine, Gly, at position 12 introduces a repulsive interaction with the phosphate group of GTP. This repulsion increases the freedom of motion for Asp, leading to increased mobility and flexibility. This increased mobility increases the space in which residues move and disrupts crucial hydrogen bonds, specifically between residues 13 and 32, and 12 and 60. The loss of these interactions destabilizes the structural order necessary for the hydrolysis of GTP to GDP.

The G12D mutation in KRAS is not only associated with structural changes, such as the disruption of hydrogen bonds and increased flexibility, but it also significantly impacts the protein’s thermodynamic properties and the network of residue interactions[10, 11]. Our analysis reveals that the mutation increases dynamic disorder or entropy within the system. As key residues become more flexible, the protein can explore a wider range of conformational states, reflected in increased entropy. Interestingly, mutual information (MI) analysis shows that while individual residues exhibit greater disorder, their interactions become more correlated and ordered. This apparent paradox of simultaneous disorder and enhanced coupling warrants further explanation.

We use the concept of “dynamic coupling,” where increased individual disorder leads to new interaction networks that paradoxically stabilize KRAS’s oncogenic behavior. Our findings establish a metric for comparing these distinct behaviors across different proteins by examining how mutations influence dynamic coupling. In contrast to our findings in KRAS, another MD study we conducted on Pyrin[12] reveals an opposite behavior: individual entropies increase less than joint entropies following mutation. This indicates that mutations in Pyrin lead to decreased interactions between residues, a phenomenon we term “dynamic decoupling.” This contrasting behavior suggests that different mutations can have fundamentally different effects on residue interactions and protein functionality. While it is evident that the G12D mutation increases the distance between Asp12 and the phosphate group, leading to increased flexibility, the precise relationship between these structural changes and observed alterations in entropy and MI remains less understood. These thermodynamic and informational metrics are crucial for understanding how the mutation disrupts finely tuned interactions necessary for KRAS’s regulatory function. Our study addresses this gap by integrating detailed MD simulations with entropy calculations and MI analysis, enhanced by Kernel Density Estimation (KDE) for greater precision. By employing KDE, we achieve a quantitative assessment of MI changes, providing deeper insights into how the mutation’s effects alter the protein’s interaction network. We propose an inequality demonstrating that changes in individual entropies of two residues i and j exceed those in joint entropy: Δ(H(i)+H(j))>Δ(H(i,j)), where H(i) and H(j) are the individual entropies and H(i,j) is the joint entropy. Using the definition of Mutual Information, this inequality is written as Δ*MI* (*i*; *j*) > 0, i.e., a positive shift in mutual information shows dynamics coupling. Through this integrated approach, we seek to explain the G12D mutation’s role in promoting persistent activation of KRAS and its implications for cancer progression. Our study highlights the value of combining multiple computational techniques, entropy analysis, MI, and KDE, to clarify atomic interactions responsible for dysfunctional regulation observed in mutant KRAS.

It is important to emphasize that the concept of dynamic coupling we use is complementary to significant advancements made in this field, notably by Ozkan et al., [13–15] who analyze the coupling between residues through perturbation-response relationships. In contrast, our description of dynamic coupling relies solely on correlations observed in unperturbed proteins, focusing on how the probability distribution function of fluctuations in one residue influences the probability distribution function of another through their joint probability distribution, thereby highlighting the way the two residues communicate. There have been studies investigating the change of hydrolysis rate upon G12D mutation using different computational tools, such as QM/MM free energy calculations[4, 16–18] and molecular dynamics[8, 9, 19–21]. Following the pioneering work of McClendon et. al[22] and LeVine et.al[23] on the use of mutual information in protein physics, there has been a large body of work investigating the impact of mutations on changes in mutual information—evaluated through atomic fluctuations derived from molecular dynamics simulations[24–29]. The present study uses mutual information and entropy analysis to understand the impact of the G12D mutation on the function of KRAS.

## Materials and methods

### i. Structure of KRAS

The structure of the wild type protein used in our study is 6GOD.pdb[30]. The ribbon diagram of the molecule is shown in Figure 1. The G12D mutant that we also used is the structure[30] 6GOF.pdb. Both of the structures have 172 residues and contain the GTP equivalent GNP as the ligand.Switch I, SI, typically involves residues 25-40. It is involved in binding to effector proteins and changes conformation when GTP or GDP is bound. Switch II, SII, typically involves residues 60-74. This region undergoes conformational changes that are crucial for the catalytic activity of the protein, particularly in the hydrolysis of GTP to GDP. These conformational changes allow KRAS to transition between active (GTP-bound) and inactive (GDP-bound) states, modulating downstream signaling pathways. The P-loop, phosphate-binding loop is formed by residues 10-17. It stabilizes the phosphate chain of GTP, especially the β- and γ-phosphates[31]. The numbering of atom numbers for GDP that we use in the figures in the results section are defined in the Supplementary Material section.

**Figure 1.**
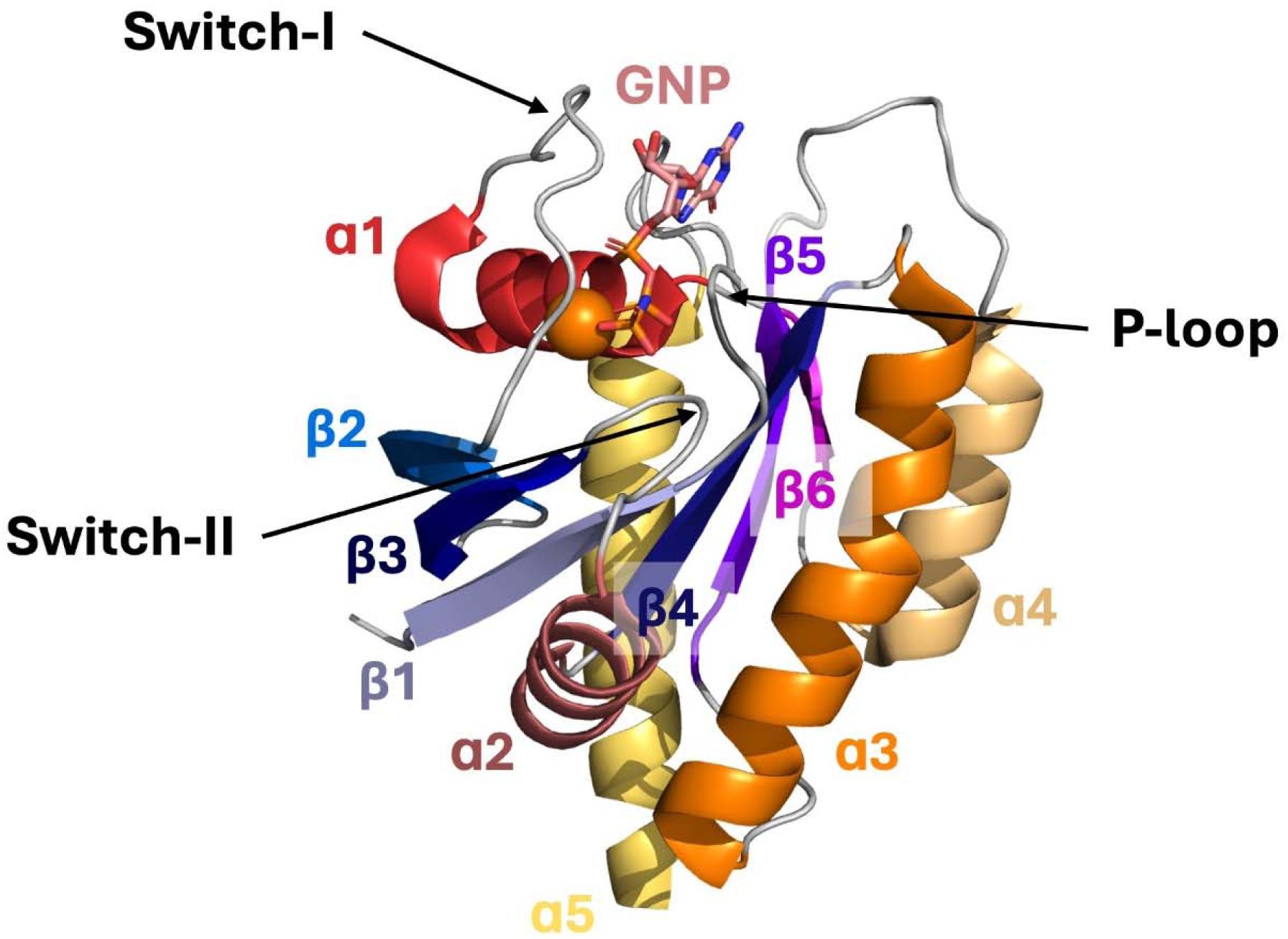
Crytal structure of KRAS showing the six beta sheets and five helices, the P-loop, Switch –I and Switch –II. The GTP equivalent of GNP is also shown in stick format.

KRAS has a β-sheet structure composed of six beta strands forming a central β-sheet flanked by alpha helices. The beta strands in KRAS are labeled as β1 to β6. In addition to providing a stable scaffold for the protein, the set of six beta strands act as a conduit of information transfer between GNP and downstream molecules such as the P-loop, Switch I, and Switch II. KRAS contains five alpha helices (α-helices), which play critical roles in its structural integrity, GTP/GDP binding, and interactions with other proteins. The alpha helices in KRAS are labeled α1 to α5, and they serve as functional components that support the protein’s dynamic movements and interactions with effectors and regulators. Among these, α5, residues 151-165, positioned near the C-terminal end is involved in membrane association and plays a role in anchoring KRAS to the plasma membrane[31].

The P-loop, Switch I, and Switch II regions transmit conformational changes induced by GTP binding and hydrolysis which propagate to effector-binding regions, allowing KRAS to signal its active or inactive state. The conformational changes in Switch I, especially in the effector loop, enable the interaction with downstream effectors like RAF, which then trigger further signaling cascades. The beta strands work together to ensure that the entire KRAS molecule undergoes coordinated motion in response to GTP/GDP binding, allowing for efficient signal transmission. All simulations are performed for the active form of KRAS in the presence of Guanosine 5’-[(α,β)-methyleno]triphosphate (GNP) which is a non-hydrolyzable analog of GTP (guanosine triphosphate). GNP stabilizes the protein in its GTP-bound state and prevents the hydrolysis of GTP.

### ii. Molecular dynamics simulations

The crystal structures of WT and G12D mutant KRAS with PDB id’s 6GOD and 6GOF, respectively, was used for the simulations, GNP and the crystal waters were retained from the structures. Both systems were prepared using CHARMM-GUI [32, 33]. The systems were solvated in a TIP3P water box with 15 Å padding and neutralized using NaCl, resulting in a salt concentration of 0.15 mol/L. Charmm36 force field[34] was used for parametrization. A cutoff of 12 Å was applied for the short-range, nonbonded interactions, and a cutoff of 10 Å for the smothering functions. Long-range electrostatic interactions were managed using the particle-mesh Ewald (PME) method[35]. The time step of 2 fs was used for integration. Both systems underwent 5,000 step energy minimizations. Following the minimization, systems were heated from 60 K to 300 K with a constant pressure of 1 atm using Nosé-Hoover Langevin piston[36]. Following the annealing step, systems were equilibrated for 1ns without any constraints on atoms. The temperature was maintained at 300 K using Langevin dynamics. Production runs were performed under NpT ensemble for 100 ns 6 replicas for each system, making the total simulation time 600 ns/system. All simulations were performed using NAMD 3.0b6.

The trajectory gives the instantaneous position *R_i_* (*t*) of all atoms i of the protein for time t defined at discrete time points. The instantaneous fluctuation Δ*R_i_* (*t*) of residue i at time t is obtained by subtracting the mean position *R̄*_*i*_ from the instantaneous position. In Cartesian coordinates, the fluctuation vector is written as

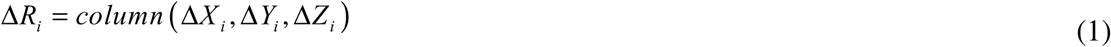

and the joint fluctuation of i and j is

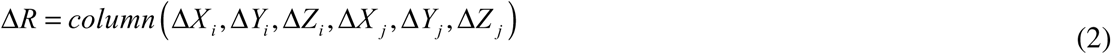

where Δ*R* constitutes the state vector defined for each time point of the MD simulations and the time argument is not shown for brevity.

### iii. Principal Component Analysis (PCA)

Principal Component Analysis (PCA) was performed to analyze the fluctuations of residues across the protein structure. The covariance matrix of the atomic displacements was computed from the simulation trajectory, and eigenvalue decomposition was applied to identify the principal components of motion. The first two principal components (PC1 and PC2), which account for the largest variance in the residue fluctuations, were visualized to assess the dominant modes of motion in the system. This approach enabled us to capture the key dynamic features of the protein, highlighting regions of significant conformational flexibility and identifying potential allosteric communication pathways.

### iv. Kernel density estimation (KDE)

Estimating quantities like entropy and mutual information from MD simulations requires modeling the probability density function (PDF) of the fluctuation vectors, Δ *R*. One option is fitting a multivariate Gaussian distribution[37], however this does not model the multimodal structure of some of the residue and residue-pair fluctuations. A more accurate estimate of the PDF can be obtained using kernel density estimation which represents the distribution as:

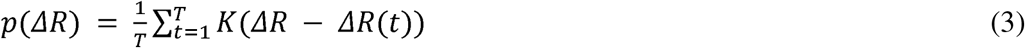

where Δ*R* is the estimated fluctuation vector for one or more residues, T is the number of simulation steps, and K is the kernel function, typically a Gaussian. For two residues i and j, for example, the estimated probability will be *p* _(_Δ*R_i_*, Δ*R_j_* _)_. KDE concentrates the distribution on the observed data points and the degree of concentration is controlled by a bandwidth parameter, typically the covariance, that determines the shape and width of the kernel. The bandwidth is tuned to maximize the likelihood of unseen data using cross-validation.

In our experiments we implemented KDE using the cuML library[38] to take advantage of GPU acceleration to rapidly calculate distributions for all residue pairs and single residues. We used isotropic multivariate Gaussians as kernels and optimized the bandwidth for each residue/residue-pair independently. Gaussians with diagonal or full covariance matrices as kernels did not significantly affect the results. We checked for the convergence of our estimation by comparing the results to those with partial data.

Convergence of trajectory lengths was assessed by comparing the cumulative mutual information values (definition provided below) for trajectories of 100 ns and 600 ns, as shown in Figure S2 of the Supplementary Material section.

### v. Entropy, Mutual information and cumulative mutual information

The Shannon entropy for a residue *i* based on the fluctuations of its Cα is defined according to

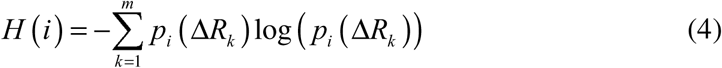

where the trajectory is divided into *m* bins according to the magnitude of the fluctuation values, and *p_i_* (Δ*R_k_*) is the probability of observing fluctuation Δ*R_k_* of residue i. We adopt log as the natural logarithm and the summation is performed over all time points along the trajectory representing the average 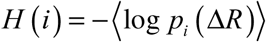. The entropy quantifies the uncertainty or disorder associated with the distribution of fluctuations of the residue. When Shannon entropy is multiplied by the Boltzmann constant *k_B_*, it can be interpreted as a statistical or information-theoretic form of thermodynamic entropy[39]. This is particularly true for systems in statistical mechanics, where the microstates of the system are considered, and their probabilities are used to calculate entropy. We use *k_B_* =1 throughout this paper. For two residues, the joint entropy similarly follows:

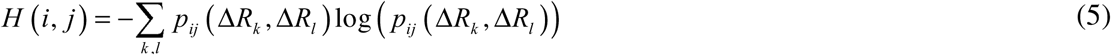

where k and l denote the bins and *p_ij_* (Δ*R_k_*, Δ*R_l_*) denotes the probability of fluctuations of i and j in the designated bins. The mutual information between a pair of residues i and j can be calculated based on the probability distributions of the fluctuations or motions of the residues. It provides a quantitative measure of the strength of the relationship between residues in terms of their dynamics, allowing one to identify residues that tend to move together or influence each other’s motions. Mathematically, MI is defined as

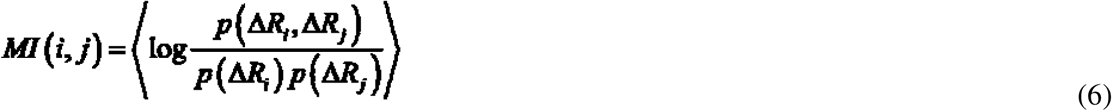

The variables are functions of time, and the angular brackets denote averaging over time. In its full generality, the joint probability in Eq. 6 is six dimensional and the singlet probability is three dimensional.

MI is written in terms of entropies as

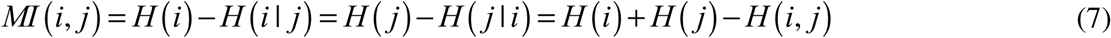

Where

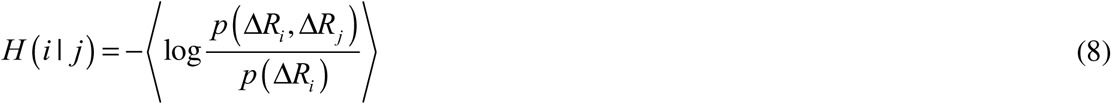

is the conditional entropy of i with respect to j.

Throughout, we obtain the probability distribution functions, PDF’s, using the non-parametric kernel density estimation, KDE, which is more accurate than the multivariate Gaussian used for probability distribution functions. It makes minimal assumptions about the underlying data distribution and can capture complex, multimodal, and non-linear dependencies. KDE can adapt to the actual shape of the data distribution, capturing more information about the interactions and dependencies between residues. It can identify subtle and complex relationships that simpler models might miss. Because it can capture more detailed and accurate dependencies, the MI values obtained are often higher compared to methods that make simplifying assumptions.

Summing MI (i, j) over the index j we obtain a measure known as the cumulative mutual information, CMI, of a residue i,

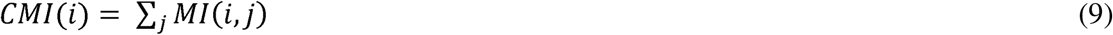

CMI profiles provide an aggregate measure of how strongly a particular residue i is interconnected with the rest of the protein. It essentially reflects the overall degree of communication or coupling between residue i and its environment within the protein structure. Higher CMI values indicate that a residue is highly embedded or coupled within the protein’s network of interactions, suggesting its significant role in the protein’s structural stability and functional dynamics. Conversely, lower CMI values may indicate residues that are less central or critical to the overall protein conformation and behavior.

We will be specifically interested in the changes in entropy, mutual information and cumulative mutual information upon mutation defined, respectively as

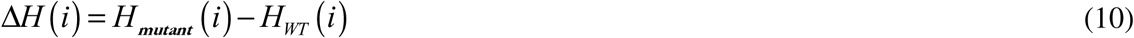

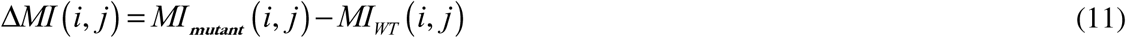

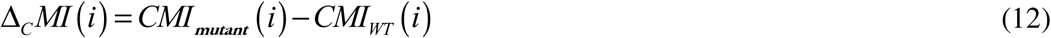

where, Δ*_C_* indicates the change in the cumulative MI. The left-hand members are the profiles when plotted as a function of the variable i.

By comparing the profile shifts, Δ*_C_MI* (*i*), for example, one can identify how mutations alter the communication pathways and interaction strengths among residues. These shifts can provide insights into the molecular basis of the mutation’s effects on protein function and stability.

In addition to the MI profile shifts shown by Eqs. 10 and 11, we will also calculate entropy profile shift upon mutation, Δ*H* (*i*) given by Eq. 10.

### vi. Flexibility

We measure the flexibility of a residue by calculating its entropy, which reflects the range of conformations the residue adopts throughout the trajectory. Thus, defining entropy based static flexibility of a residue by *f_SH_*

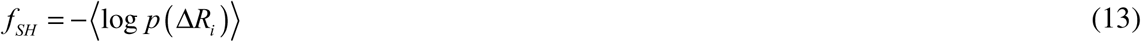

The entropy defined above by Eq.13 in terms of Cartesian coordinates can also be calculated from the internal coordinates of the protein defined by the so-called Ramachandran angles *ϕ* and *ψ*. Throughout the molecular dynamics (MD) trajectories, the frequency of passage from one state to another may also be used to define flexibility using the Ramachandran variables. Accordingly, we define a transition-based flexibility by the equation

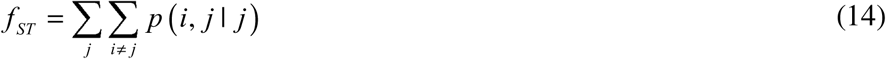

where *p*(*i*, *j* | *j*) is the conditional probability of having the *ϕ* angle in state i, given that the*ψ* angle is in state j. Since, in general, *p*(*i*, *j* | *j*) will not be equal to *p*( *j*,*i* | *i*) we take the average

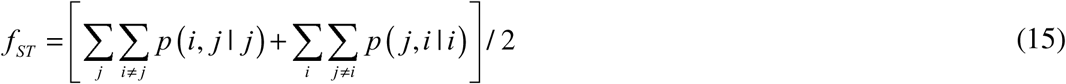

The entropy-based flexibility measures the extent of space probed by the residue, providing insights into its overall conformational freedom. Transition-based flexibility, on the other hand, reflects how the internal coordinates of the residue change, capturing more subtle shifts in its position and orientation. The changes in their values due to mutation are meaningful and provide complementary insights into the residue’s dynamic behavior. We will utilize both entropy-based and transition-based expressions in evaluating flexibilities to gain a comprehensive understanding of how mutations influence protein dynamics.

### vii. Estimating the allosteric network from MI

To estimate the allosteric network in KRAS, we employed a method combining structural proximity and mutual information (MI) analysis. First, a contact map was generated by identifying residue pairs within a cutoff distance of 7.3 Å, which reflects physical proximity between residues likely to be involved in direct interactions. Next, we calculated the MI heat map to assess the degree of correlated motion between residue pairs. A heat map is a contour map, shown in Figure S-1 in the Supplementary Material Section, presented in terms of residue indices. In this figure, contacting residues and those with values of MI greater than a given threshold are shown with a point. We considered residue pairs with MI values greater than 0.5 to have strong coupling, indicative of functional or allosteric relationships.

The intersection of these two datasets, presented as Figure S1 in the Supplementary Material Section, identifying residues that are both physically close (within the cutoff distance) and exhibit high MI, was used to define the potential allosteric paths. These residues are considered to play key roles in transmitting allosteric signals through the protein. As shown in Figure S1, this procedure highlighted all of the beta strands of the protein as the members of the allosteric network, consistent with the idea that beta sheets can act as pathways for mechanical and functional signaling in proteins. This approach provides a structural and dynamic basis for identifying key regions involved in allosteric regulation within the protein.

## Results

The analysis of our MD simulations on WT and G12D KRAS consists of three parts. In part A, we examine structural changes in inter-residue distance fluctuations, hydrogen bonds, and internal coordinates caused by the mutation. While these concepts have been explored in earlier research[5, 8, 9, 19, 40–43], our analysis extends beyond them, focusing on inter-residue distance changes to set the stage for a deeper investigation into entropy and mutual information variations. In part B, we analyze entropy and mutual information changes, which offer new insights into the dynamic coupling of KRAS residues. This aspect, not widely explored before, provides a more detailed understanding of the molecular dynamics and interactions that underpin KRAS’s behavior in both its wild-type and mutated forms. We also connect these mutual information changes to explain allosteric interactions. In part C, we discuss the dynamic interactions of KRAS with atoms of GNP and the coordination of water molecules.

### A. Dynamic changes upon mutation that lead to increase of flexibility

The crystal structures of KRAS, specifically 6GOD (WT) and 6GOF (G12D mutant), show that the distance of 4.37Å between the Cα of G12 and the γ-phosphate of GNP is larger than the corresponding distance of 4.29Å in WT. This increased distance is fundamentally driven by the repulsion between the negatively charged aspartic acid at position 12 and the negatively charged phosphate group of GNP. This repulsion lies at the core of all subsequent changes that lead to the malfunctioning of KRAS. As noted earlier, the fluctuations in the distance between residue 12 and the phosphate group become more pronounced. The results of molecular dynamics trajectories lead to a PDF for distances shown in Figure 2, clearly illustrate the dynamic basis of this critical difference. In WT, shown by the solid line in the figure, the distance distribution has a sharp peak while the mutant shown with the dot-dashed curve is spread out, extending to larger distances. The increase in the fluctuations has far-reaching consequences, which we will analyze in depth in this section.

**Figure 2.**
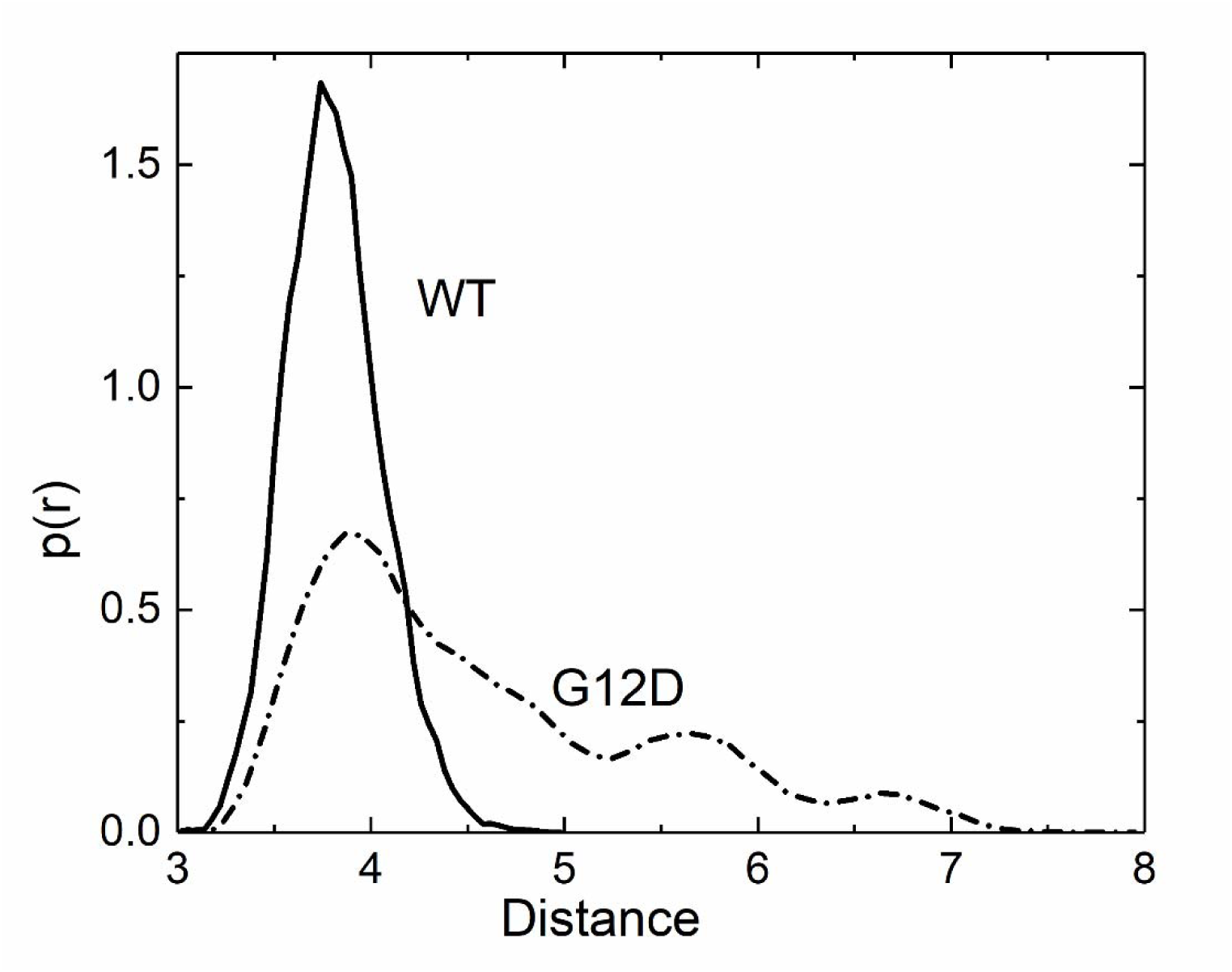
The distance distribution of the Cα of residue 12 and the γ-phosphate of GNP in W (solid line) and G12D mutant (dashed line) cases. The curves are obtained by using the Multiple Peak Fit module of the software Origin[44] that gives the best fit through the set of data points. To avoid cluttering the figure, the points derived from binning the molecular dynamics trajectories are not displayed.

#### a. Mutation dependence of distance fluctuations

The immediate consequence of the increase of the G12D-GNP fluctuations is the distribution of the G12-Y32 distance shown in Figure 3. The WT has a sharp peak similar to the one in Figure 2. The mutant shows two peaks one of which is above 12 Å resulting from interactions of D12, GNP and Y32. As a result, the Switch1 loop that contains Y32 can take a larger number of conformations.

**Figure 3.**
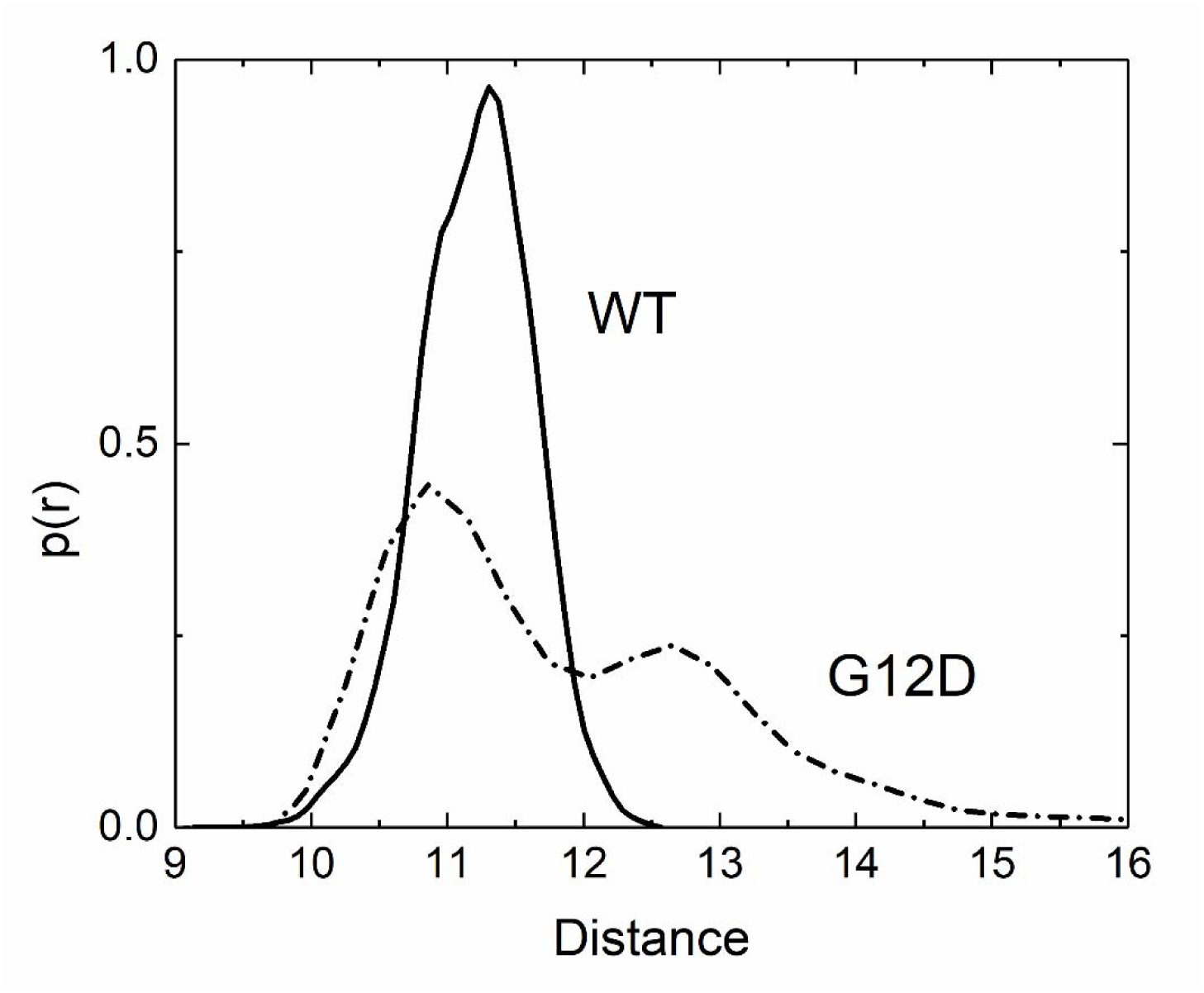
The distance distribution of the Cα of residue 12 and Y32 in WT (solid line) and G12D mutant (dashed line) cases. See legend to Figure 2.

The distances between alpha carbons of residues 12 and 61 in the crystal structure are 14.33Å and 14.36Å in WT and G12D, respectively. Molecular dynamics shows a more dramatic difference as shown by the two curves in Figure 4. Each has a double peak, the smaller peak corresponding to the crystal structure distances and the second peak from inter-residue interactions during the evolution of the dynamics. Both peaks are at larger distances in the mutant, making Q61 more mobile, which affects the ability of Q61 in its role in the hydrolysis of GNP.

**Figure 4.**
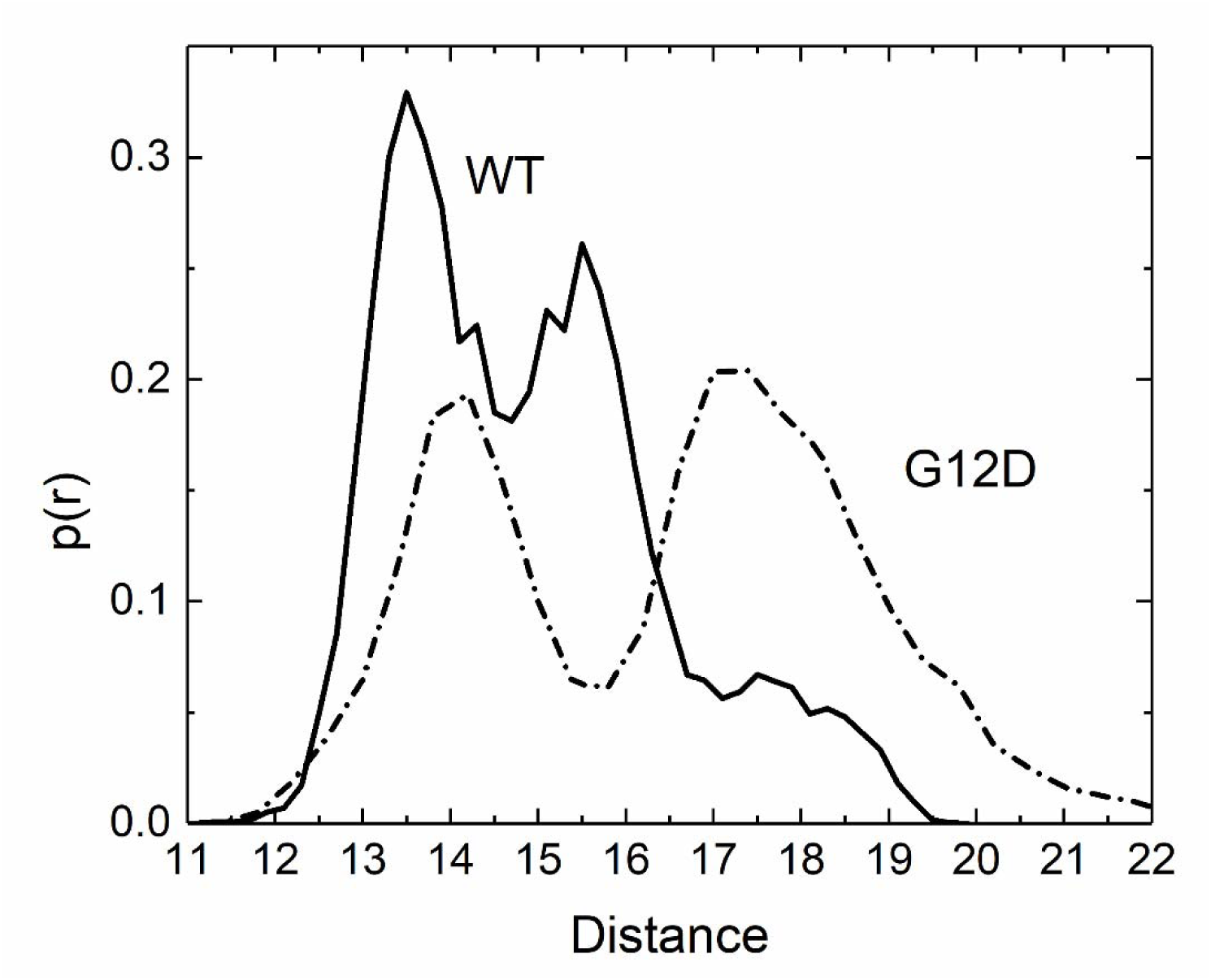
The distance distribution of the Cα of residue Y32 and Q61 in WT (solid line) and G12D mutant (dashed line) cases. See legend to Figure 2.

#### b. The G13-Y32 and T35-G60 hydrogen bonds are broken in the G12D mutant trajectories

Molecular dynamics simulations show that the G12D mutation dramatically reduces the frequency of this critical hydrogen bond—from 70% in the wild-type to only 3% in the mutant. This disruption not only breaks a key structural constraint but also leads to a marked increase in entropy, reflecting substantial changes in the protein’s dynamic behavior. The G13-Y32 hydrogen bond is crucial for maintaining the proper conformation of the P-loop and Switch I regions, which are essential for sequestering GNP within active KRAS. The rupture of this bond causes a structural rearrangement, allowing these regions to become more flexible and adaptive, enabling them to engage more closely with GNP. This structural shift enhances the protein’s ability to interact more intimately with GNP (see below), stabilizing these interactions and promoting stronger binding affinity. This intimate interaction is observed by the increased mutual information (MI) both within the residue pairs of the protein and between the protein and GNP. We refer to this increased MI as ‘dynamic coupling’. Trajectories also show a hydrogen bond between T35 and G60 which has a frequency of 100% for the WT and 38% for the mutant. The hydrogen bond parameters are selected as 5 Å and 40 degrees for both cases in VMD *Hbonds* plugin.

#### c. Introduction of multiple peaks on the Ramachandran maps

Analysis of the WT and G12D crystal structures shows that there is a consistent increase in the distances between Cα pairs in the mutant, while this consistent trend is not seen in the side chain atom distances. Such distance changes are well determined by torsion angles □ and ψ, whose fluctuations represent the statistics of the protein backbone. The variance of these angles is key to the protein’s ability to adopt various conformations, which can be critical for binding to other molecules, undergoing allosteric changes, or performing its biological function. The splitting of a single □-ψ peak into two peaks, as observed in the G12D mutation, indicates increased flexibility and shows that the mutation has introduced a new stable conformation. This splitting also triggers the splitting of several other □-ψ peaks along the loop and the neighboring loops, in the form of an allosteric effect, allowing the protein to adopt new conformations that are more favorable for GNP binding. This allosteric shift explains the observed increase in flexibility, as the protein explores a wider range of conformational states to optimize GNP interaction.

The splitting of the □-ψ peak for Y32 is shown in Figure 5. When we compare the left and right panels in Figure 5, we can see that Y32 can adopt multiple conformations upon G12D mutation, whereas in the WT case it’s restricted to a single conformation.

**Figure 5.**
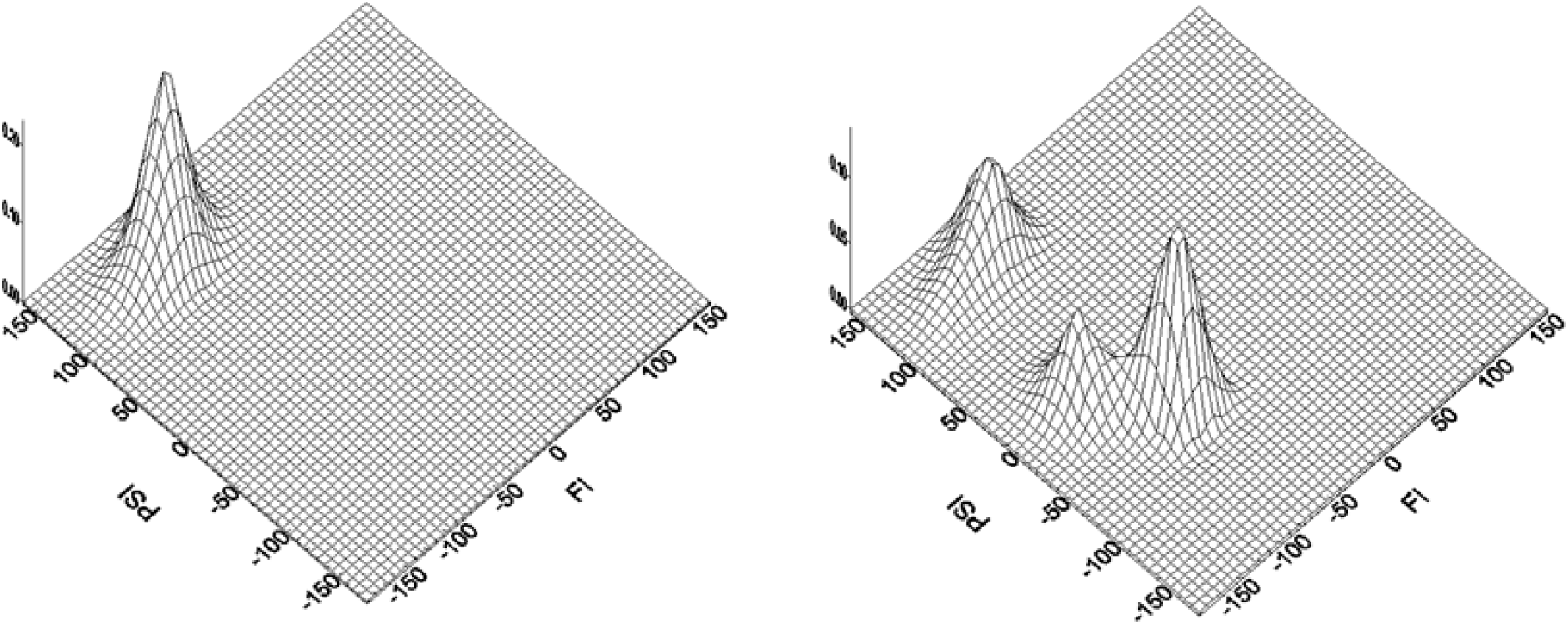
The Ramachandran plot of Y32 in WT (left panel) and G12D (right panel) cases. Several other residues exhibit peak splitting. These residues are located on Switch I and II. The figures for some of these are presented in the Supplementary Material section, Figures S3 and S4.

#### d) Changes in Root Mean Square Deviation (RMSD) of residues

The changes in the fluctuations of residues are shown in Figure 6. Residue indices between 1 and 172 are for the protein, those larger than 172 refer to the atoms of GNP. Atom numbers for GNP are identified in the Supplementary Material section. The largest increase in RMSD values is for Switch 1, with the largest value for residue I36. To a lesser extent the residues of Switch 2 show increased RMSD in the mutant with largest values for G60 and S65. The largest peak on the right hand side of the figure is for residues M170, S171 and K172. The increased RMSDs in these C-terminal residues, despite their distance from the binding site and GNP, indicate longer-range communication within the protein structure resulting from changes in the protein’s flexibility or dynamic equilibrium, leading to structural adjustments even in distant regions. The C-terminal of KRAS is known to interact with membranes and can play roles in signaling and localization. Changes in this region, as observed in the increased RMSD for these residues, could suggest alterations in membrane interactions or other distal regulatory mechanisms.

**Figure 6.**
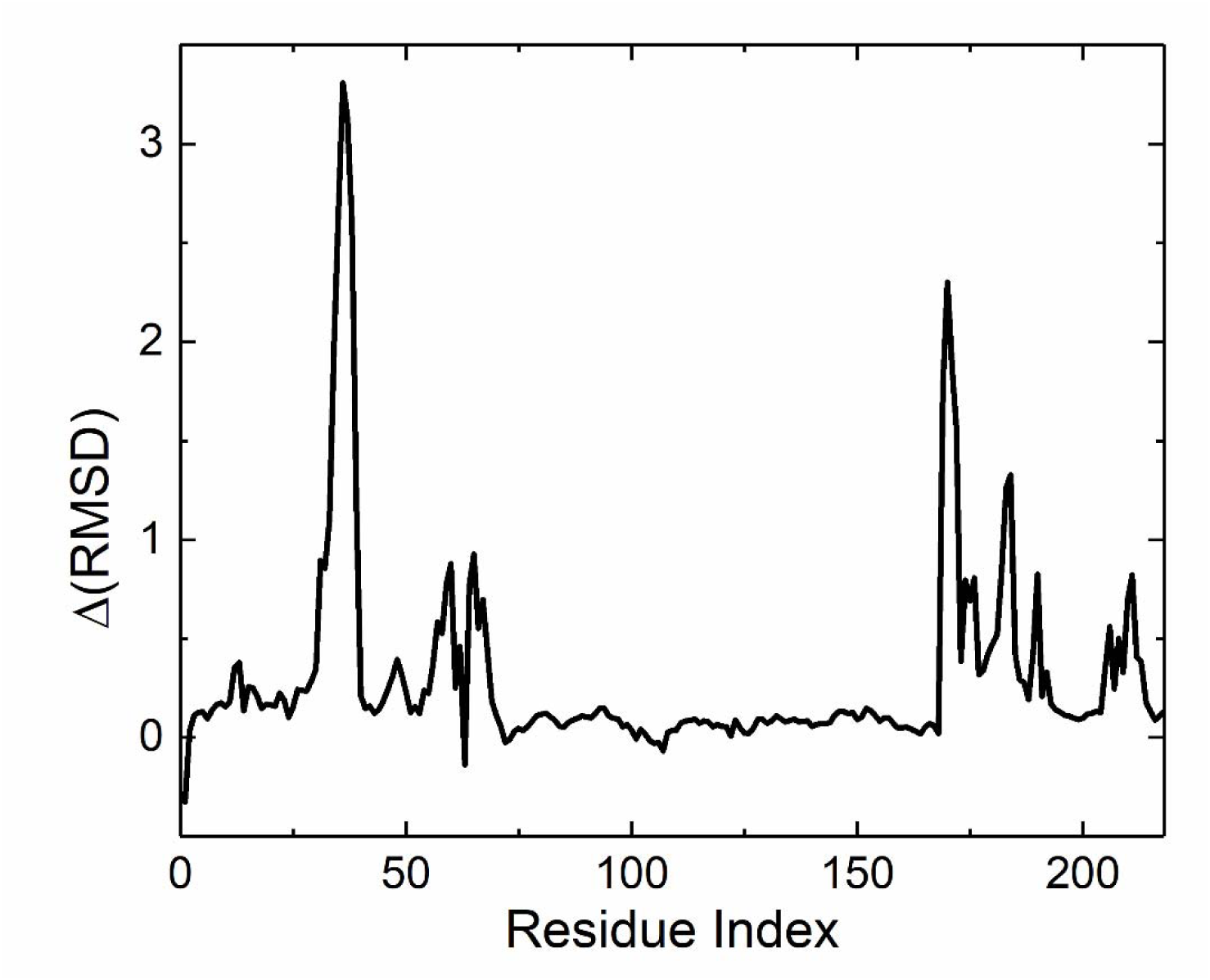
The change of RMSDs (RMSD_G12D_-RMSD_WT_) of KRAS and GNP. Residues from 1 to 172 belong to KRAS and those larger than 172 refer to the atoms of GNP.

#### e) Residues 12, 32, 60 and 61 and their effects on the hydrolysis of GNP

These residues play significant role on the function of KRAS. They are part of a coordinated mechanism where several key residues contribute to the correct positioning of the water molecule that attacks the γ-phosphate of GNP, leading to its hydrolysis.

The fluctuation distributions of these residues in WT and G12D structures are presented in Figures 7.

**Figure 7.**
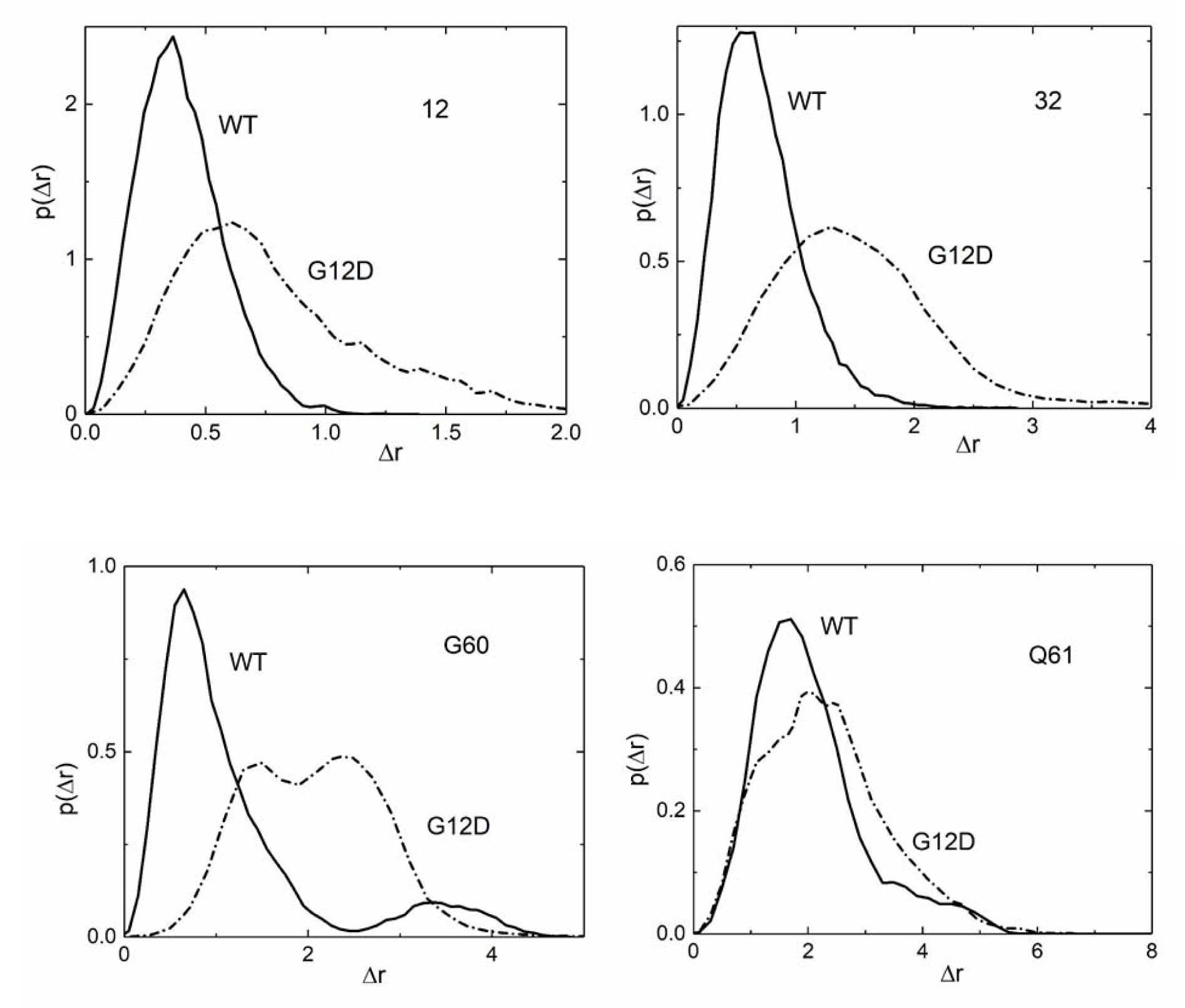
The fluctuation distribution function *p*(Δ*r*) of residues 12(upper left panel), Y32(upper right panel), G60(lower left panel) and Q61(lower right panel), where solid lines are for WT and dashed lines are for G12D cases.

Following the observed increases in fluctuations of residues 12, 32, 60, and 61 in the G12D mutant, we can conclude that these changes disrupt critical functions. G12, essential for the tight binding and proper orientation of the γ-phosphate of GNP, loses its ability to maintain control due to increased distance and flexibility. Similarly, Y32 in Switch I, which stabilizes the γ-phosphate through hydrogen bonding, experiences increased fluctuations, reducing its regulatory control. The high mobility of G60 and Q61, both involved in positioning a water molecule for nucleophilic attack on the γ-phosphate, disrupts the precise geometry needed for efficient GTP hydrolysis. This loss of structural coordination significantly impairs the hydrolysis process in the mutant.

In order to understand the dynamic interactions of KRAS with GNP further, we analyzed the interactions of the protein with GNP throughout the WT and G12D simulations and detected the major changes of hydrogen bonding upon G12D mutation. We found the hydrogen bonds in both simulations by using the *Hbonds* plugin of VMD with a distance cutoff of 3.5 Å and angle cutoff of 30°. We only considered the hydrogen bonds with an occupancy higher than 10% of the simulation time. In Table 1. We show how the occupancies of hydrogen bonds change upon mutation. Upon close inspection we detected major changes, decreased bond occupancies. The residues that lose hydrogen bond interactions with GNP are 12, 13, 15, 16,18 and 32.

**Table 1.**
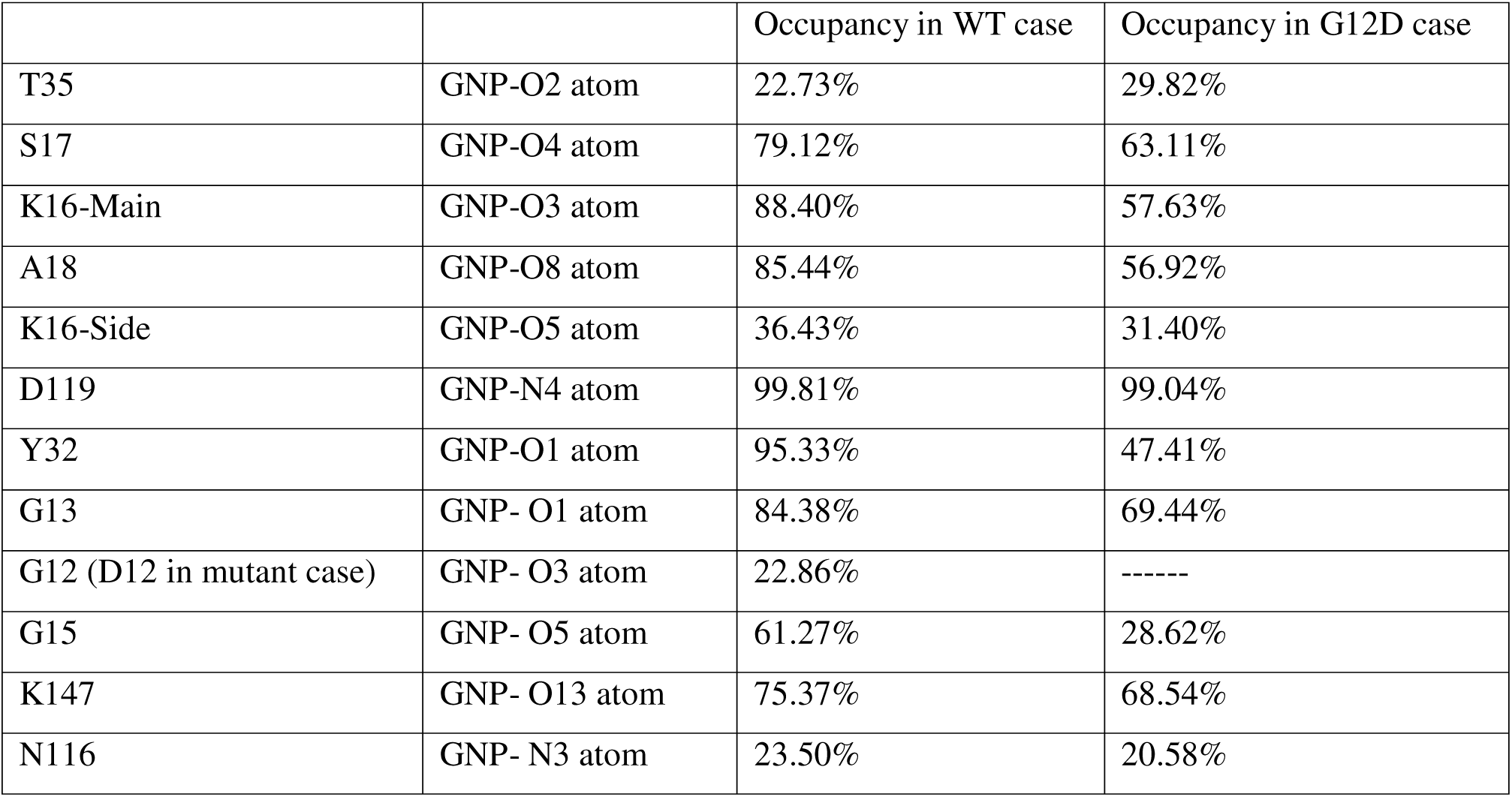
Hydrogen bonds with occupancies higher than 10% for WT and G12D KRAS.

|  |  | Occupancy in WT case | Occupancy in G12D case |
| --- | --- | --- | --- |
| T35 | GNP-O2 atom | 22.73% | 29.82% |
| S17 | GNP-O4 atom | 79.12% | 63.11% |
| K16-Main | GNP-O3 atom | 88.40% | 57.63% |
| A18 | GNP-O8 atom | 85.44% | 56.92% |
| K16-Side | GNP-O5 atom | 36.43% | 31.40% |
| D119 | GNP-N4 atom | 99.81% | 99.04% |
| Y32 | GNP-O1 atom | 95.33% | 47.41% |
| G13 | GNP- O1 atom | 84.38% | 69.44% |
| G12 (D12 in mutant case) | GNP- O3 atom | 22.86% | ----- |
| G15 | GNP- O5 atom | 61.27% | 28.62% |
| K147 | GNP- O13 atom | 75.37% | 68.54% |
| N116 | GNP- N3 atom | 23.50% | 20.58% |

We performed a Principal Component Analysis (PCA) on the fluctuations of these residues and compared their fluctuation profiles in WT and G12D simulations. Figure 8 shows the comparison of PCA’s of WT and G12D cases for selected residues and the comparison of total number of hydrogen bonds of the protein with GNP. From Figure 8 panels A, E and F, we can conclude that these residues. 12 18 and 32, became more flexible and have more conformational states.

**Figure 8.**
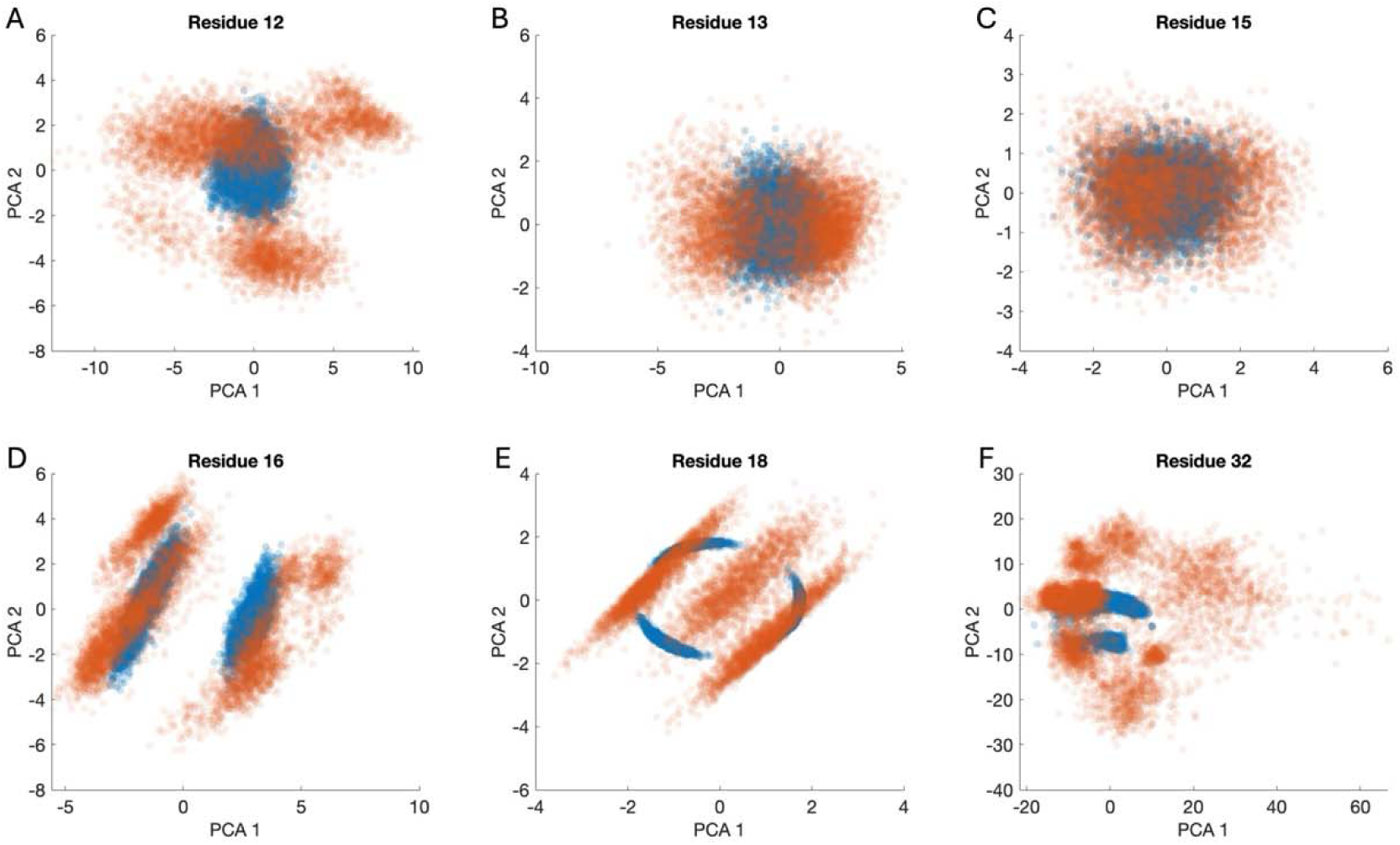
PCA of residue fluctuations of residues 12, 13, 15, 16, 18 and 32 are shown in panels a to F, where blue is for WT fluctuations and orange is for G12D mutation.

### B. Entropy and Mutual Information Changes

#### a) Mutation Increases Entropies of Residues

In Figure 9. we present the entropy increase profile resulting from the mutation and relate these changes to the fluctuations and interactions of important residues.

**Figure 9.**
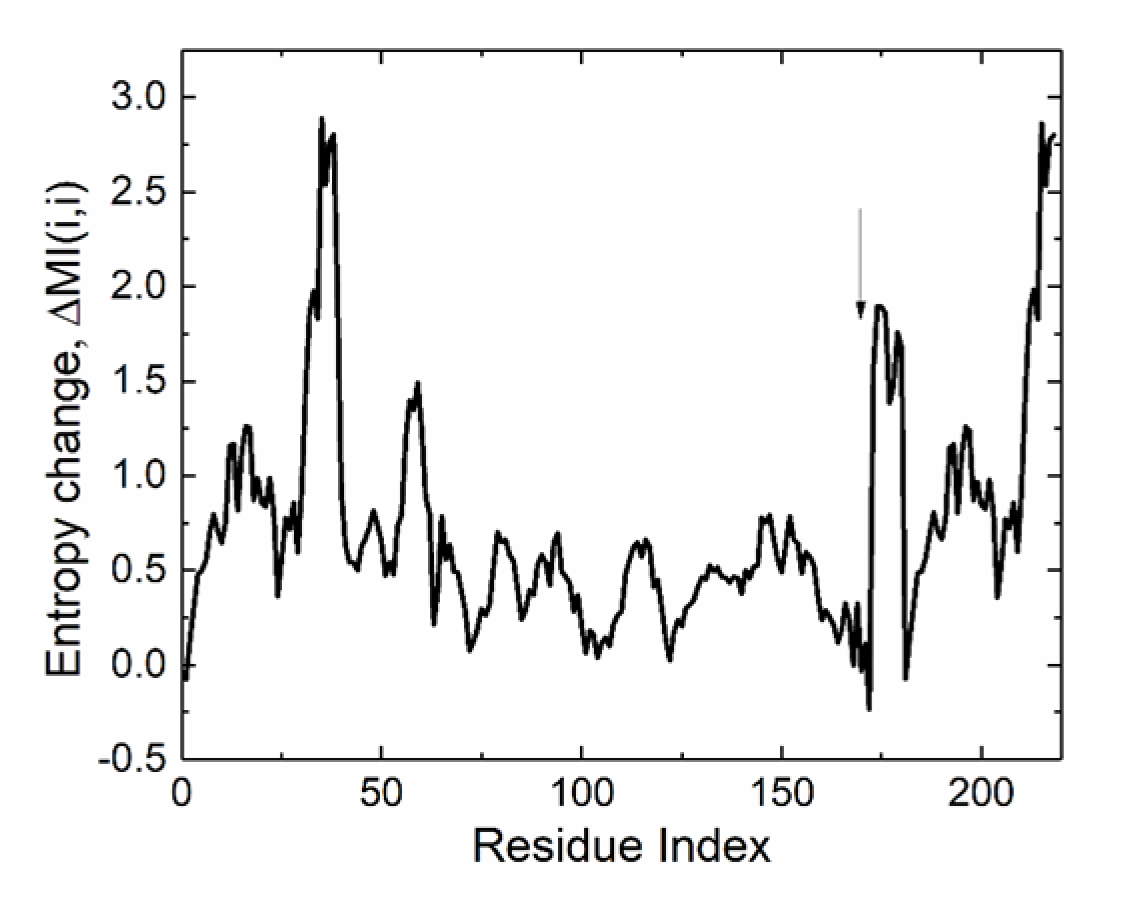
The difference of residue entropies of G12D from WT case for KRAS and GNP. Residues from 1 to 172 belong to KRAS and those larger than 172 refer to the atoms of GNP. The G12D mutation leads to significant increases in the entropies of specific residues, particularly D12, Y32, G60, and Q61, along with the phosphate groups and tail atoms of GNP. These entropy increases correspond to enhanced flexibility and disruption of crucial interactions, as described below:

The increase in the entropy of residue 12 results from the mutation of glycine to aspartic acid, which introduces a larger, negatively charged side chain. This additional bulk and charge lead to greater flexibility and mobility, increasing the distance between D12 and the γ-phosphate of GNP. The repulsion destabilizes critical interactions that position GNP, reducing the control of residue 12 over the hydrolysis process. As may be seen from Table 1, the increase in entropy is significant going from 0.07 to 1.23. Transition based flexibility, on the other hand, is small and increases from 0.0082 to 0.0093, showing that G12 probes different regions in space but does not change its internal coordinates much.

The increase in the entropy of Y32 results from the breaking of the hydrogen bond with G13 and the splitting of the available Ramachandran angles. Its entropy and transition-based flexibilities go from 1.52 to 2.89 and this increase in flexibility disrupts Y32’s ability to stabilize the GTP-binding pocket through its interaction with residue 12. The greater mobility of Y32 misaligns the catalytic components, contributing to inefficient hydrolysis.

The increase in the entropy of G60 results from heightened fluctuations after the mutation. In the wild type, G60 plays a role in coordinating water molecules for nucleophilic attack on the γ-phosphate. The increased flexibility in the mutant disrupts the precise positioning of these water molecules, impeding the proper geometry needed for hydrolysis. This change may be seen from the increase of flexibility from Table 2. Entropy based flexibility goes from 1.87 to 3.12. Transition based flexibility goes from 0.108 to 0.805. The Ramachandran angles for G60 are shown in Supplementary Material Figure S4 which shows that in G12D residue G60 gains freedom of almost fully unhindered Glycine.

**Table 2.** Flexibility Changes upon G12D Mutation

|  | Entropy |  | Transition Based Flexibility |  |
| --- | --- | --- | --- | --- |
| | WT Entropy | G12D Entropy | WT $f_{ST}$ | G12D $f_{ST}$ |
| G12/D12 | 0.07 | 1.23 | 0.0082 | 0.0093 |
| Y32 | 1.52 | 2.89 | 0.015 | 0.036 |
| G60 | 1.87 | 3.12 | 0.108 | 0.805 |
| Q61 | 3.21 | 4.11 | 0.187 | 0.806 |

The increase in the entropy of Q61 results in a reduced ability to coordinate the two water molecules involved in GTP hydrolysis. In the mutant, Q61 fluctuates more and binds only a single water molecule, which does not participate in catalysis. This miscoordination further disrupts the hydrolysis process by preventing the proper positioning of the catalytic water molecule. The changes in flexibility are significant and similar to those of G60.

The increase in entropy is also observed in the phosphate groups and tail atoms of GNP, indicating greater flexibility in these regions. This could contribute to the destabilization of the nucleotide-binding pocket, further interfering with GTP hydrolysis.

#### b) The G12D mutation leads to an overall increase in mutual information (MI) between residue pairs

This change is visualized in the heatmap of Figure 10, where red points indicate a decrease in MI, with values ranging from −0.2 to −0.1. Gray points signify an increase, with values between 0.5 and 1.0, while black dots mark the strongest increases, ranging from 1.0 to 5.8.

**Figure 10.**
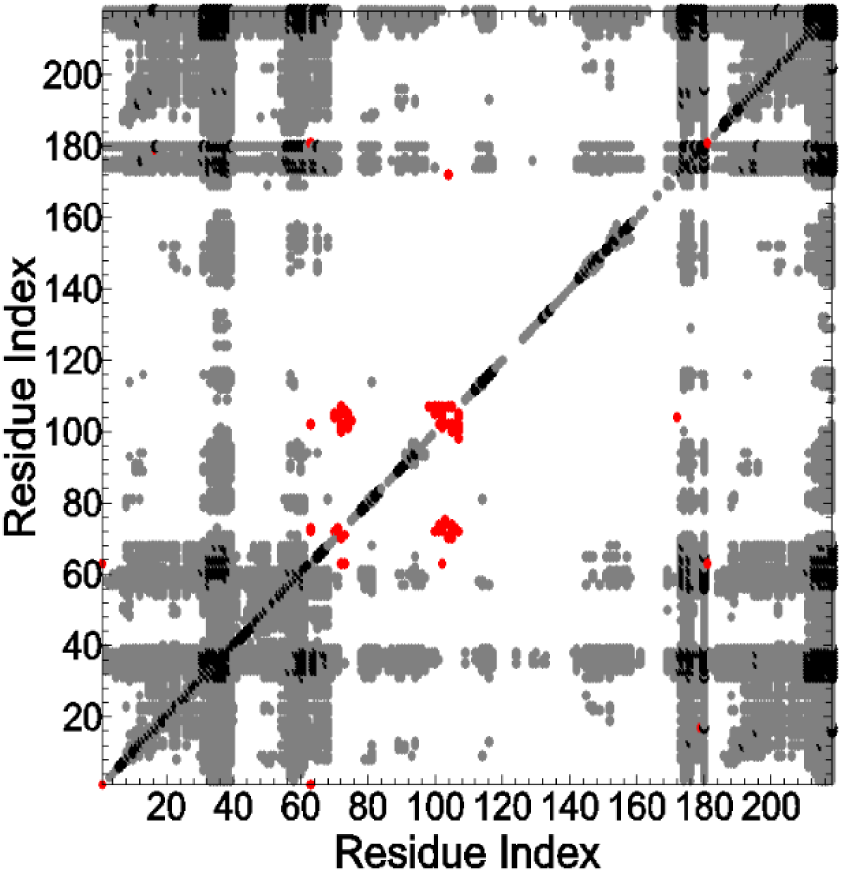
Difference of Mutual Information (MI) between residue pairs upon G12D mutation. Red dots indicate a decrease in MI, values between −0.2 to −0.1, whereas gray and black dots indicate an increase of MI between 0.5 and 1.0 and 1.0 to 5.8, respectively,

Using Eqs 10-11 the positive shift condition Δ*MI* (*i*, *j*) observed from Figure 10 is written in terms of entropy changes as

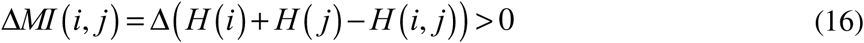

As discussed in the preceding sections, entropy H(i) of residue i reflects the uncertainty or disorder associated with the fluctuations of that residue. A higher entropy means that the residue fluctuates more and is less predictable. Joint entropy H(i,j) reflects the combined uncertainty or disorder of two residues, i and j, considered together. If the two residues are strongly correlated, their joint entropy will be lower than the sum of their individual entropies, because knowing the state of one residue gives information about the state of the other.

Upon mutation, residues i and j both experience an increase in their individual entropies (Figure 9). This means that each residue’s behavior becomes more uncertain or disordered. This increase in individual entropy suggests that the mutation increases the flexibility or mobility of these residues, allowing them to explore a larger number of conformational states.

The joint entropy of the two residues, H(i,j), increases as well, but not as much as the sum of the individual entropies as may be deduced from Eq. 16 and the positive values of MI. This suggests that, although each residue is more disordered individually, they retain some level of correlation or interaction with each other. We examine the consequences of this dynamic coupling in detail in the discussion section.

The gray streaks in the heat map represent residues with specific functional roles that show increased correlations with several other residues in KRAS due to the G12D mutation. These vertical and horizontal bands highlight regions that are not only functionally important but also dynamically more connected in the mutant.

Band 1: range 30-36, part of Switch I, which plays a crucial role in nucleotide binding and the regulation of KRAS activity. Specifically, Y32, D33, P34, S35 and G38 show increased coupling with the rest of the protein. In the G12D mutant, this region’s increased mobility (or increased MI with other residues) suggests altered dynamics that may disrupt KRAS’s normal function, contributing to its malfunction in regulating GTP hydrolysis. The formation of new hydrogen bonds, changes in flexibility, and interaction patterns all impact the switch I region’s ability to properly regulate the active/inactive states of KRAS.

Among these residues, T35 shows the strongest interactions with the rest of the protein and GNP. In Figure 11, we show the MI shift values for T35 and remaining residues and GNP atoms. The figure shows that the coupling of T35 with Q61 and S65 increases strongly, both of which are outside the first coordination shell of T35. The coupling of T35 increases also with the phosphate and tail atoms of GNP.

**Figure 11.**
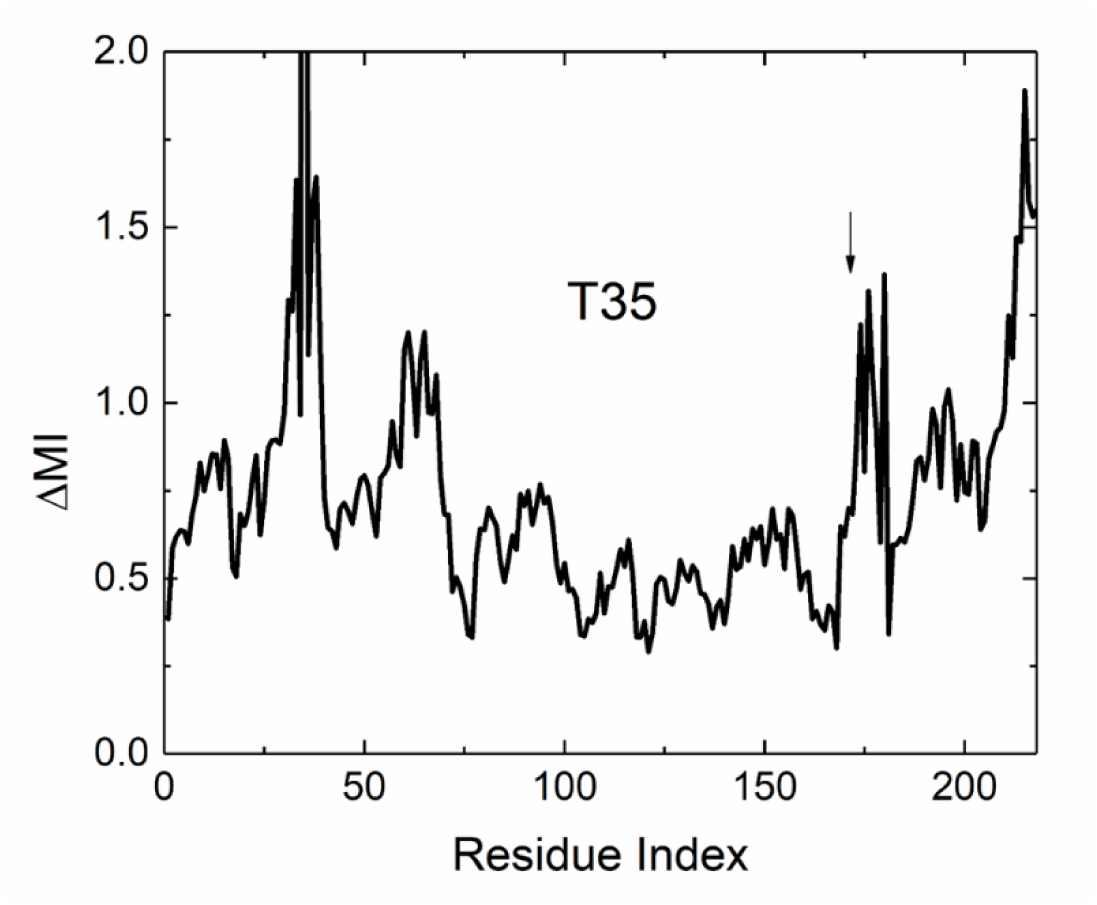
The change of MI of T35 upon G12D mutation with the other residues in KRAS and GNP. Residues from 1 to 172 belong to KRAS and those larger than 172 refer to the atoms of GNP. The arrow shows the onset of GNP atoms.

Band 2, range 58-65 of Switch II, which is essential for the regulation of KRAS function, particularly in GTP hydrolysis and signal transduction. Specifically, D57, G60, Q61, E62 and Y64 show increased coupling. Increased mobility in these residues indicates a loss of structural rigidity that is required for effective GTP hydrolysis. The changes in MI or fluctuations within this band reflect a destabilized and dysfunctional Switch II region in the mutant form. In Figure 12 we show the MI shifts of Q61.

**Figure 12.**
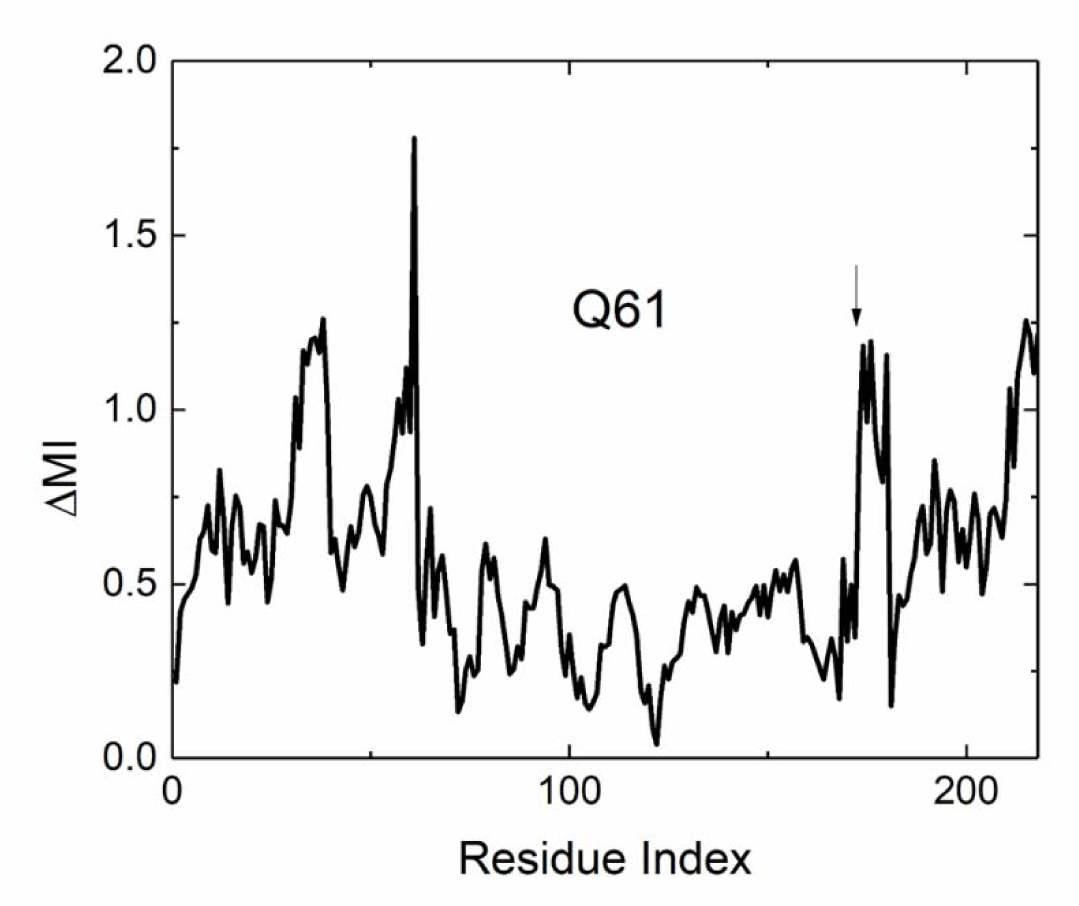
The change of MI of Q61 upon G12D mutation with the other residues in KRAS and GNP. Residues from 1 to 172 belong to KRAS and those larger than 172 refer to the atoms of GNP.

Band 3, range 170-172 in the C-terminal region is involved in the membrane association of the protein. Specifically, M170 is part of the α5 helix, which plays a role in positioning KRAS relative to the membrane and facilitating its interaction with other proteins in signal transduction pathways. Residues 171-172 are in the hypervariable region, HVR, just after the α5 helix. The HVR is crucial for post-translational modifications, such as farnesylation and palmitoylation, which help anchor KRAS to the cell membrane. These residues are less involved in the catalytic function of KRAS but are critical for its correct localization to the plasma membrane. In Figure 13, we show the MI shifts of residue K172.

**Figure 13.**
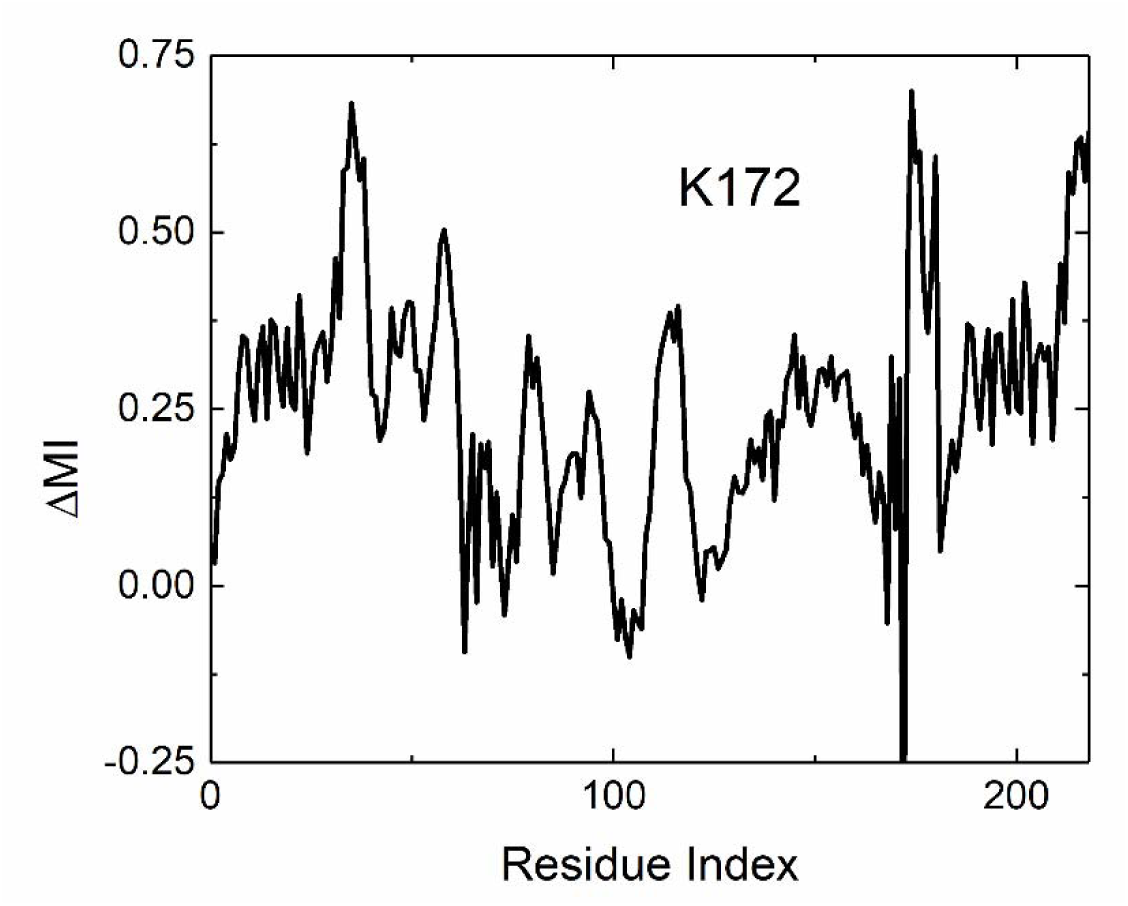
The change of MI of K172 upon G12D mutation with the other residues in KRAS and GNP. Residues from 1 to 172 belong to KRAS and those larger than 172 refer to the atoms of GNP.

The increased fluctuations or RMSD values, Figure 6, and increased coupling in the mutant, Figure 9, suggest that the mutation indirectly affects KRAS’s membrane-binding properties, potentially disrupting its proper orientation or interaction with other signaling molecules. This could interfere with KRAS’s ability to transmit signals from the membrane to downstream effectors. While it may initially seem coincidental, the fact that several distant residues show increased coupling in the mutant suggests a systematic change **i**n how the protein dynamics have been altered by the G12D mutation. The mutual information increase between these regions is likely indicative of a real dynamic coupling, which reflects long-range communication or coupled motions that emerge as a response to the structural and functional changes induced by the mutation.

Q61, T35, and residues 170-172 all exhibit an increase in MI with GNP, indicating that their motions are more tightly coupled to GNP in the mutant. This increased MI reflects enhanced correlated motion, suggesting that these residues interact with GNP more dynamically compared to the wild type. The G12D mutation disrupts the typical coordination of residues and water molecules involved in hydrolysis, leading to compensatory dynamic interactions between these residues and GNP.

The increased MI between residues 170-172 and GNP is particularly striking, as these residues are distant from the binding site and not traditionally involved in hydrolysis. This implies that the G12D mutation not only affects the local environment but also leads to long-range allosteric effects which indicates that the mutation may be affecting the overall dynamics and flexibility of the protein in a way that extends beyond the active site, contributing to global changes in the

#### c) Weak interactions in KRAS

We also present Figure 14, which highlights the regions with weak interactions, identified by negative MI shifts. This information may be crucial for drug design strategies. By targeting these weak interaction regions, it may be possible to develop drugs that counteract the effects of the G12D mutation by stabilizing these regions or inhibiting their altered interactions. Such an approach could potentially oppose the mutation’s effects, offering a targeted therapeutic strategy.

**Figure 14.**
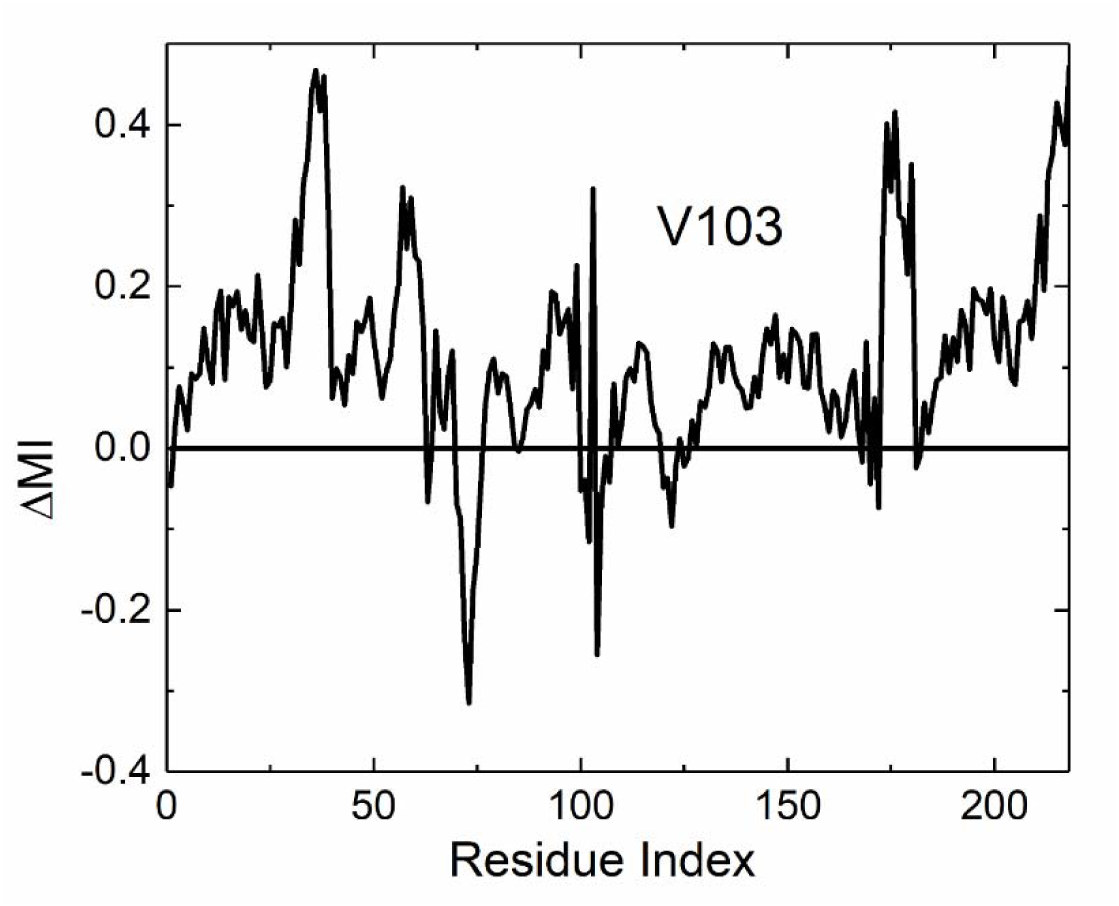
The change of MI of V103 upon G12D mutation with the other residues in KRAS and GNP. Residues from 1 to 172 belong to KRAS and those larger than 172 refer to the atoms of GNP.

#### d) Cumulative MI profile shifts

In Figure 15 we present the cumulative MI shift plot for the protein and GNP. The profile shows several peaks corresponding to residues that gained importance as the result of mutation. The residues corresponding to the peaks in the CMI profile are likely to be dynamically coupled to many other residues in the protein. A high CMI value for a residue indicates that it shares significant mutual information with multiple other residues, suggesting that these residues are involved in coordinated movements or interactions during protein dynamics. The peaks indicate key communication pathways within the protein.

**Figure 15.**
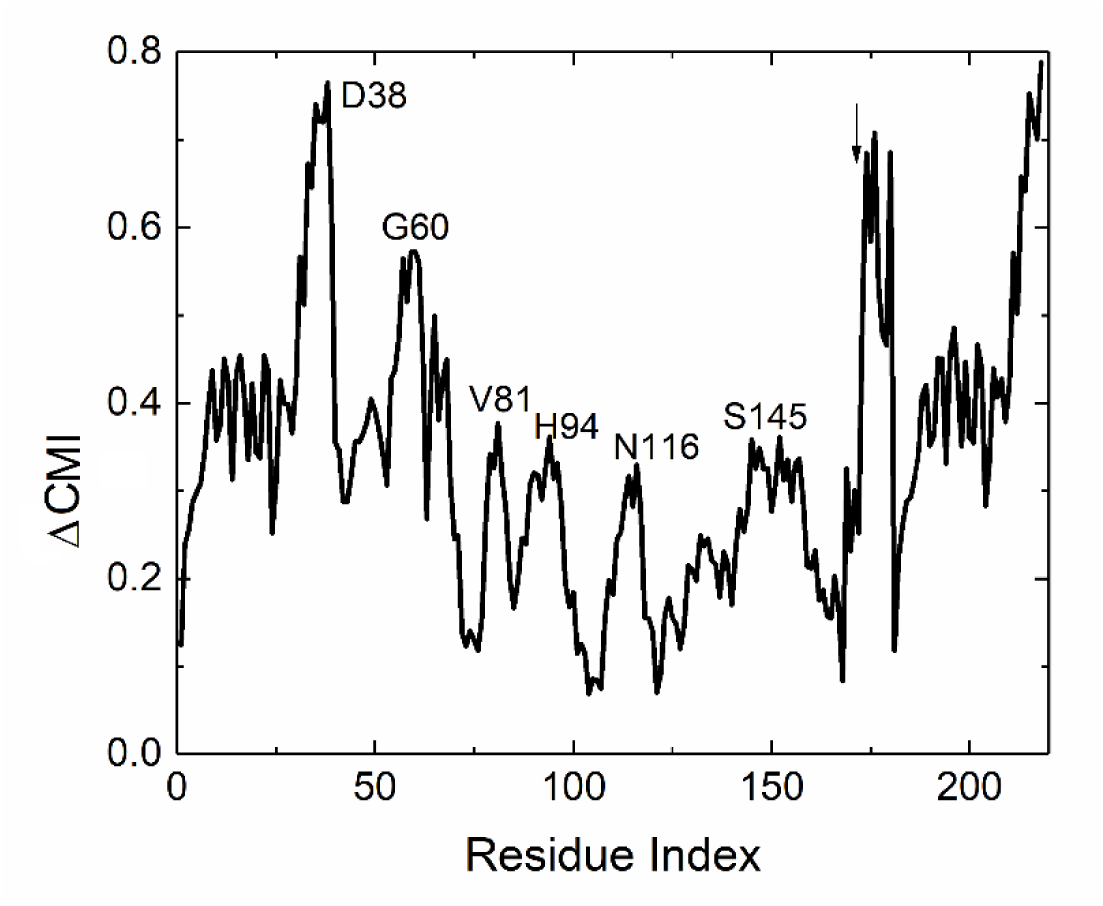
Cumulative MI shift values. The important peak residues are indicated on the plot. The starting point of GNP is shown with the small arrow in the figure.

H94 is on helix4 and G60 is at the terminus of a beta strand. The remaining four peaks are on beta strands which are predicted as residues on allosteric paths (See Section C below).

### C. Dynamics of interaction of KRAS with atoms of GNP and the coordination of bridging water molecules

In our MD simulations, we identified two stable bridging water molecules located between KRAS residues and the GNP. These water molecules play a key role in stabilizing the coordination of the Mg² ion. By comparing their behavior in the WT KRAS and the G12D mutant simulations, we observed notable changes in their interactions with nearby residues. We focused on hydrogen bonds with occupancies greater than 10%, ensuring that only significant interactions were considered.

Upon comparison of Table 3 (WT) and Table 4 (G12D), we found that the overall occupancy of hydrogen bonds involving the water molecules and residues D33 and T58 decreased in G12D, suggesting a weakening of these interactions. However, the interaction with D57 remained stable in both simulations. Interestingly, the G12D mutation led to the formation of a new hydrogen bond between the water molecules and residue T57, which was absent in the WT case. We observed the average number of bridging water molecules between the protein and GNP molecule. For the WT on average there are 7.16 water molecules and for G12D on average there are 7.48 water molecules within the 5 Å shell around the GNP binding cavity. The G12D mutation decreases the occupancy of some hydrogen bonding between the protein and the water molecules but it creates new ones and the negatively charged aspartate in G12D variant enhances the interactions with water molecules and Mg², altering the hydration network. This can explain the increase in the MI values between the protein and GNP, indicating that the G12D mutation not only alters the local hydration network but also enhances the dynamic coupling between the protein and GNP. These changes in water-mediated interactions and hydrogen bond dynamics likely contribute to the stabilization of the G12D mutation, influencing the overall structural and functional behavior of KRAS in its GTP-bound state.

**Table 3.** Hydrogen bond occupancies from WT simulations, between the bridging water molecules and KRAS.

| Donor | Acceptor | Occupancy |
| --- | --- | --- |
| TIP3 - 332-Side | ASP57-Side | 97.44% |
| TIP3 - 326-Side | ASP33-Main | 85.12% |
| TIP3 - 332-Side | THR58-Main | 84.29% |

**Table 4.** Hydrogen bond occupancies from G12D simulations, between the bridging water molecules and KRAS.

| Donor | Acceptor | Occupancy |
| --- | --- | --- |
| TIP3 - 329-Side | ASP33-Main | 35.95% |
| TIP3 - 328-Side | ASP57-Side | 82.34% |
| TIP3 - 328-Side | THR58-Main | 55.41% |
| TIP3 - 329-Side | THR35-Side | 22.93% |
| TIP3 - 329-Side | ASP57-Side | 13.76% |
| TIP3 - 328-Side | THR35-Side | 15.00% |

\*TIP3 – 326 from WT simulations is equivalent to TIP3 – 329 In G12D simulations, and TIP3 – 332 from WT simulations is equivalent to TIP3 – 328 in G12D simulations.

Figures showing MI shifts between KRAS and GNP atoms are presented in the Supplementary
Material section.

### C. Allostery in KRAS

Two distinct types of allosteric effects have been identified in KRAS[45, 46]. The first involves local allosteric changes centered around residue 12, which influence the interactions of specific residues in the Switch I (Y32, D33, T35) and Switch II (Q61, E62, Y64) regions, modulating effector binding. The second involves long-range allosteric effects that originate from residue 12 and extend to residues 168-172, affecting KRAS’s interactions with membranes. These pathways highlight the dual role of G12 as an allosteric hub, with both functional and spatial consequences for KRAS activity.

#### 1. Local Allosteric Effects on Effector Binding

The first type of allosteric regulation we observe in KRAS is a local effect, where the G12D mutation influences the protein’s interaction with downstream effectors. Effector proteins, such as RAF kinases, bind to KRAS at the switch regions, which are known to be directly impacted[4] by the structural changes induced by the mutation. The G12D mutation alters the local conformational landscape, particularly in residues like G12, Y32, and Q61, leading to increased flexibility and dynamic disorder.

This localized increase in flexibility disrupts the precise interaction network required for proper effector binding. Specifically, the mutation affects the communication between these key residues, leading to a “local allosteric effect”. Despite the mutation’s effect being confined to the spatially close residues, its impact on effector binding has significant downstream consequences[4], as it alters KRAS’s ability to transmit signals correctly. Although we describe this as a “local allosteric effect”, due to its initiation at spatially close residues, the distances between G12 and Y32 (9.94 Å) and between Y32 and Q61 (14.33 Å) indicate that these residues lie outside the first coordination sphere. As a result, the disruption of key residue interactions significantly impacts effector binding and the overall signaling function of KRAS, emphasizing the global consequences of a localized perturbation.

#### 2. Distant Allosteric Effects Involving C-Terminal Residues E168-K172 and Membrane Interactions

The second type of allostery we observe involves long-range effects that extend beyond the local mutation site, involving residue 172 in KRAS. Our analysis shows that the increase of MI between the mutation site and residues E168-K172 plays a crucial role in distant allosteric regulation by influencing KRAS’s interaction with the membrane.

Unlike local allostery, which is confined to the switch regions, the allosteric effect involving residue K172 suggests that KRAS can transmit information from its GTP-binding site (where mutations like G12D occur) to distant parts of the protein. This long-range coupling affects the orientation and membrane-binding behavior of KRAS, likely impacting its interaction with other membrane-bound proteins or downstream partners. The dynamic coupling between residues G12, Y32, and Q61, together with residues E168-K172, enables KRAS to modulate its membrane interaction, which is critical for its biological function.

Interestingly, this distant allosteric pathway is distinct from the mutation’s direct effects on the effector-binding interface. Instead, the interaction through C-terminal residues with the membrane highlights how the G12D mutation may also influence KRAS’s spatial positioning and mobility within the membrane, affecting its accessibility to effectors and other regulatory proteins. This form of long-range allosteric communication shows KRAS’s ability to integrate local mutational changes and propagate them across the protein to modulate membrane interactions, contributing to its oncogenic behavior.

The observation that several peaks in the cumulative mutual information (MI) profile shift curve, Figure 15, specifically at E49, S65, V81, N116, and S145, lie on beta sheets is particularly notable. Beta sheets are known to act as structural conduits for long-range allosteric communication within proteins[47, 48]. In the case of KRAS, this suggests that the G12D mutation significantly alters allosteric signaling pathways that rely on these beta strands. The beta sheets in KRAS may facilitate the propagation of conformational changes from the site of mutation (G12) to distant regions of the protein, thereby affecting the dynamic interactions of KRAS with other molecules. This finding highlights the crucial role beta sheets play not only in maintaining the protein’s architecture but also in transmitting the dynamic coupling that is essential for KRAS’s biological function. The perturbation of these regions suggests that the mutation has reprogrammed the allosteric network, potentially influencing how KRAS interacts with effector molecules and membranes. The involvement of these residues in beta strands shows the structural importance of beta sheets as “communication wires”, mediating the mutation’s effects across the protein’s functional landscape, ultimately contributing to the disruption of KRAS’s signaling pathways.

## Discussion

In this study, we have explored the impact of mutations on the entropic relationships between residues in a protein, specifically focusing on how these changes can improve our understanding of allosteric communication and functional dynamics. By analyzing the expression Δ(H(i)+H(j))−ΔH(i,j), we quantify the differences between the changes in individual entropies of residues and their joint entropy, allowing us to understand how mutations affect their interactions.

The following discussion outlines three distinct cases derived from this expression, each showing different residue coupling and dynamics as a result of mutation.

The quantity Δ(H(i)+H(j)) −ΔH(i,j), which is the change in mutual information between the i j pair due to mutation, measures the difference between the change in the sum of the individual entropies of residues i and j and the change in their joint entropy. We adopted this expression to understand the impact of the mutation on mutual information between these residues. It is instructive to discuss three cases of this equation in relation to the effects of mutation:

Case 1: ΔMI(i,j)=0 or Δ(H(i)+H(j)) −ΔH(i,j)=0

This case indicates that the increase (or decrease) in the sum of individual entropies of residues i and j is exactly matched by the increase (or decrease) in their joint entropy. This suggests that (i) The mutation does not introduce any new constraints or correlations between residues i and j. Their entropies change independently of each other, and (ii) the mutation does not significantly alter the interactions between these residues. They might still function relatively independently, with little change in their coupled dynamics. As an example, if residues i and j are located far from each other or are part of separate structural or functional regions, the mutation may not affect their relationship. For instance, two loop residues not involved in direct interactions may have independent entropic responses to the mutation.

Case 2: ΔMI(i,j)<0 or Δ(H(i)+H(j)) <ΔH(i,j)

In this case, which we call “dynamic decoupling”, the increase in the joint entropy, ΔH(i,j), is larger than the sum of the increases in individual entropies, ΔH(i)+ΔH(j). This suggests that the mutation has caused a substantial increase in the overall uncertainty of the joint system of residues i and j, beyond what would be expected from their individual contributions. The larger increase in joint entropy means that (i) the combined system of residues i and j is exploring a more disordered or diverse set of states, which indicates that the mutation has introduced significant changes to the way these two residues interact, leading to an overall increase in the system’s uncertainty, or (ii) the joint entropy has increased more than the sum of individual entropies, suggesting that residues i and j have lost some of their mutual constraints or coupling. In the wild type, these residues may have been more tightly coordinated or coupled, but the mutation disrupts this, allowing them to behave more independently and explore a wider range of conformations together.

Case 2 could indicate a loss of functional coordination between residues that were previously interacting or constrained. As an example, if residues i and j were part of a rigid structural element, like a beta-strand or a helix, but the mutation disrupted this structure. Now, instead of moving together as a part of a single element, they move more independently, leading to a large increase in joint entropy.

Case 3: ΔMI(i,j)>0 or Δ(H(i)+H(j))>ΔH(i,j)

In this case, which we call “dynamic coupling”, the increase in the sum of individual entropies, Δ(H(i)+H(j)), is greater than the increase in joint entropy, ΔH(i,j). This suggests that the mutation has led to an increase in individual disorder for residues i and j, but their joint uncertainty decreases.

This suggests that (i) the individual entropies of residues i and j have increased, indicating that each residue has gained more freedom or is exploring a broader set of conformations independently. However, the smaller increase in joint entropy suggests that their collective behavior is still somewhat coordinated or constrained, indicating a stronger mutual dependence than in the wild type, or (ii) the mutation may have introduced changes that increase the flexibility of each residue independently, but the residues still exhibit a high degree of mutual correlation. This could happen if the mutation affects the local environment of each residue without disrupting their direct interaction, or it could even strengthen their coupling. As a result, even though they are more flexible individually, they still “move” together in a correlated manner, or (iii) the mutation might induce a compensatory mechanism where, despite increased individual fluctuations, the residues adapt in such a way that their joint fluctuations remain relatively constrained. This could be due to new interactions forming between them or through their involvement in a shared structural or functional unit that restricts their joint behavior.

The present analysis shows that KRAS is a Case 3 protein. The G12D mutation introduces significant alterations in the protein’s flexibility, as evidenced by changes in entropy and mutual information between residue pairs. These changes reflect an enhanced dynamic coupling, where residues engage in more correlated motions due to the increased flexibility. This concept of dynamic coupling is central to understanding how structural changes induced by the mutation disrupt normal KRAS function, particularly in the context of GTP hydrolysis. Upon mutation, key residues such as G12, Y32, G60, Q61, and others exhibit significant increases in individual entropy, reflecting heightened flexibility. This increase in entropy leads to a broader conformational landscape for the protein, which in turn enables more intimate and dynamic interactions between these residues. The increase in mutual information, particularly between residues like T35 and GNP, readily seen in Figure 11, highlights stronger dynamic coupling between regions that are spatially distant yet functionally significant. The MI changes are not limited to direct interactions but reflect broader, coordinated motions across the protein. C-terminal residues such as E168-K172, despite being away from the GTP-binding site, exhibit increased MI with both T35 and GNP, suggesting that the increased flexibility allows distant regions of the protein to influence critical functional sites more strongly. At the same time, changes in water coordination observed in the simulations offer additional insights into how the G12D mutation affects the dynamic behavior of KRAS. The two stable bridging water molecules that coordinate the Mg² ion between KRAS residues and GNP are particularly noteworthy. Upon G12D mutation, alterations in the hydrogen bonding network, such as the decreased occupancy between water molecules and residues D33 and T58, alongside the formation of a new interaction with T57, suggest a shift in the local water-mediated interactions.

Moreover, the fact that water molecules form new interactions upon mutation might point to compensatory mechanisms that help retain some level of stability. These water-mediated interactions could buffer the effects of increased flexibility in KRAS, ensuring that critical functional sites remain properly aligned for GTP hydrolysis. Therefore, while residues become more dynamically coupled through enhanced flexibility, the water molecules provide additional stability, helping to regulate the balance between flexibility and coordinated motion.

Another important observation is the splitting of peaks in Ramachandran plots, indicative of the altered conformational states adopted by the protein post-mutation. This peak splitting suggests the emergence of new dynamics, where residues can undergo transition between different conformational states more easily, contributing to the overall flexibility of the protein.

While barrier lowering could theoretically play a role in these dynamics by affecting transition rates between states, its impact on mutual information remains unclear. MI reflects correlations in residue movements, rather than transition kinetics, so any effects of barrier lowering would be more relevant to transition rates and the decay of autocorrelations, rather than MI itself. Thus, barrier lowering, though potentially important, requires further investigation before its role in dynamic coupling can be confirmed.

In conclusion, the G12D mutation leads to increased flexibility, as evidenced by entropy changes, which in turn enables more extensive dynamic coupling between residues. This dynamic coupling, reflected in the MI increases, plays a significant role in disrupting the normal function of KRAS by altering the interactions required for GTP hydrolysis.

Traditional biochemical and structural studies often focus on static snapshots of protein structures and specific interactions, such as binding sites or mutation-induced changes in conformation. While this is valuable, proteins like KRAS are highly dynamic systems where function is influenced not just by static structures but also by continuous fluctuations and molecular motions. Entropy quantifies the degree of these fluctuations and provides a measure of the flexibility and disorder within the protein. This gives us insights into how a mutation, like G12D, increases the freedom of motion of residues, altering the protein’s behavior and function in ways that static structural data might miss. Mutual Information captures correlations between the motions of different parts of the protein. Even if two residues are far apart, MI helps us see if their motions are dynamically coupled, indicating functional or relationships. This is crucial for understanding allosteric regulation or long-range effects of mutations like G12D, where the mutation site influences distant functional residues.

By incorporating entropy and MI, we get a systems-level perspective on KRAS function. Instead of focusing on isolated interactions (e.g., single hydrogen bonds or binding events), we can analyze how the mutation impacts the entire network of residue interactions across the protein. This is particularly useful for studying proteins like KRAS, where distant regions communicate dynamically.

Drug design traditionally aims to block or enhance specific interactions. However, through entropy and MI analysis, we can see how mutations like G12D disrupt the normal flexibility and coordination within the protein. This opens the door to designing drugs that don’t just block a specific interaction but help restore normal dynamics—“re-tuning” the protein to behave more like the wild type. Such drugs would not only inhibit the pathological function of mutant KRAS but also restore the healthy, functional dynamics of the protein.

## References

[1] Zhu C, Guan X, Zhang X, Luan X, Song Z, Cheng X, et al. Targeting KRAS mutant cancers: from druggable therapy to drug resistance. Molecular Cancer. 2022;21:159.

[2] Gorfe AA, Grant BJ, McCammon JA. Mapping the nucleotide and isoform-dependent structural and dynamical features of Ras proteins. Structure. 2008;16:885–96.

[3] Lukman S, Grant BJ, Gorfe AA, Grant GH, McCammon JA. The distinct conformational dynamics of K-Ras and H-Ras A59G. PLoS computational biology. 2010;6:e1000922.

[4] Pantsar T. The current understanding of KRAS protein structure and dynamics. Computational and structural biotechnology journal. 2020;18:189–98.

[5] Chen J, Wang L, Wang W, Sun H, Pang L, Bao H. Conformational transformation of switch domains in GDP/K-Ras induced by G13 mutants: An investigation through Gaussian accelerated molecular dynamics simulations and principal component analysis. Computers in Biology and Medicine. 2021;135:104639.

[6] Chen J, Zhang S, Zeng Q, Wang W, Zhang Q, Liu X. Free energy profiles relating with conformational transition of the switch domains induced by G12 mutations in GTP-bound KRAS. Frontiers in Molecular Biosciences. 2022;9:912518.

[7] Grudzien P, Jang H, Leschinsky N, Nussinov R, Gaponenko V. Conformational dynamics allows sampling of an “active-like” state by oncogenic K-Ras-GDP. Journal of Molecular Biology. 2022;434:167695.

[8] Vatansever S, Erman B, Gümüş ZH. Oncogenic G12D mutation alters local conformations and dynamics of K-Ras. Scientific reports. 2019;9:11730.

[9] Vatansever S, Erman B, Gümüş ZH. Comparative effects of oncogenic mutations G12C, G12V, G13D, and Q61H on local conformations and dynamics of K-Ras. Computational and structural biotechnology journal. 2020;18:1000–11.

[10] Buhrman G, Casey O, Zerbe B, Kearney BM, Napoleon R, Kovrigina EA, et al. Analysis of binding site hot spots on the surface of Ras GTPase. Journal of molecular biology. 2011;413:773–89.

[11] Grant BJ, Lukman S, Hocker HJ, Sayyah J, Brown JH, McCammon JA, et al. Novel allosteric sites on Ras for lead generation. PloS one. 2011;6:e25711.

[12] Hacisuleyman A, Gul A, Erman B. Role of Mutual Information Profile Shifts in Assessing the Pathogenicity of Mutations on Protein Functions: The case of Pyrin Mutations. bioRxiv. 2024:2024.04.07.588500.

[13] Ose NJ, Butler BM, Kumar A, Kazan IC, Sanderford M, Kumar S, et al. Dynamic coupling of residues within proteins as a mechanistic foundation of many enigmatic pathogenic missense variants. PLoS computational biology. 2022;18:e1010006.

[14] Campitelli P, Modi T, Kumar S, Ozkan SB. The role of conformational dynamics and allostery in modulating protein evolution. Annual review of biophysics. 2020;49:267–88.

[15] Modi T, Ozkan SB. Mutations utilize dynamic allostery to confer resistance in TEM-1 β-lactamase. International journal of molecular sciences. 2018;19:3808.

[16] Berta D, Gehrke S, Nyíri K, Vértessy BG, Rosta E. Mechanism-Based Redesign of GAP to Activate Oncogenic Ras. Journal of the American Chemical Society. 2023;145:20302–10.

[17] Menyhárd DK, Pálfy G, Orgován Z, Vida I, Keserű GM, Perczel A. Structural impact of GTP binding on downstream KRAS signaling. Chemical science. 2020;11:9272–89.

[18] Calixto AR, Moreira C, Kamerlin SCL. Recent advances in understanding biological GTP hydrolysis through molecular simulation. ACS omega. 2020;5:4380–5.

[19] Mir SA, Nayak B, Aljarba NH, Kumarasamy V, Subramaniyan V, Dhara B. Exploring KRas Protein Dynamics: An Integrated Molecular Dynamics Analysis of KRas Wild and Mutant Variants. ACS omega. 2024;9:30665–74.

[20] Chen C-C, Er T-K, Liu Y-Y, Hwang J-K, Barrio MJ, Rodrigo M, et al. Computational analysis of KRAS mutations: implications for different effects on the KRAS p. G12D and p. G13D mutations. PloS one. 2013;8:e55793.

[21] Shi S, Zheng L, Ren Y, Wang Z. Impacts of mutations in the P-loop on conformational alterations of KRAS investigated with Gaussian accelerated molecular dynamics simulations. Molecules. 2023;28:2886.

[22] McClendon CL, Friedland G, Mobley DL, Amirkhani H, Jacobson MP. Quantifying correlations between allosteric sites in thermodynamic ensembles. J Chem Theory Comput. 2009;5:2486–502.

[23] LeVine MV, Perez-Aguilar JM, Weinstein H. N-body information theory (NbIT) analysis of rigid-body dynamics in intracellular loop 2 of the 5-HT2A receptor. arXiv preprint arXiv:14064730. 2014.

[24] Cortina GA, Kasson PM. Excess positional mutual information predicts both local and allosteric mutations affecting beta lactamase drug resistance. Bioinformatics. 2016;32:3420–7.

[25] Erman B. Mutual information analysis of mutation, nonlinearity, and triple interactions in proteins. Proteins: Structure, Function, and Bioinformatics. 2023;91:121–33.

[26] Miyashita N, Yonezawa Y. Mutual information analysis of the dynamic correlation between side chains in proteins. The Journal of Chemical Physics. 2021;155.

[27] Hacisuleyman A, Erman B. Synergy and anti-cooperativity in allostery: Molecular dynamics study of WT and oncogenic KRAS-RGL1. Proteins: Structure, Function, and Bioinformatics. 2024;92:665–78.

[28] Habgood M, Seiferth D, Zaki A-M, Alibay I, Biggin PC. Atomistic mechanisms of human TRPA1 activation by electrophile irritants through molecular dynamics simulation and mutual information analysis. Scientific Reports. 2022;12:4929.

[29] Verkhivker GM. Biophysical simulations and structure-based modeling of residue interaction networks in the tumor suppressor proteins reveal functional role of cancer mutation hotspots in molecular communication. Biochimica et Biophysica Acta (BBA)-General Subjects. 2019;1863:210–25.

[30] Cruz-Migoni A, Canning P, Quevedo CE, Bataille CJ, Bery N, Miller A, et al. Structure-based development of new RAS-effector inhibitors from a combination of active and inactive RAS-binding compounds. Proceedings of the National Academy of Sciences. 2019;116:2545–50.

[31] Simanshu DK, Nissley DV, McCormick F. RAS proteins and their regulators in human disease. Cell. 2017;170:17–33.

[32] Jo S, Kim T, Iyer VG, Im W. CHARMM-GUI: a web-based graphical user interface for CHARMM. Journal of computational chemistry. 2008;29:1859–65.

[33] Jorgensen WL, Chandrasekhar J, Madura JD, Impey RW, Klein ML. Comparison of simple potential functions for simulating liquid water. The Journal of chemical physics. 1983;79:926–35.

[34] Lee J, Cheng X, Jo S, MacKerell AD, Klauda JB, Im W. CHARMM-GUI input generator for NAMD, GROMACS, AMBER, OpenMM, and CHARMM/OpenMM simulations using the CHARMM36 additive force field. Biophysical journal. 2016;110:641a.

[35] Darden T, York D, Pedersen L. Particle mesh Ewald: An N⋅ log (N) method for Ewald sums in large systems. The Journal of chemical physics. 1993;98:10089–92.

[36] Martyna GJ, Tobias DJ, Klein ML. Constant pressure molecular dynamics algorithms. The Journal of chemical physics. 1994;101:4177–89.

[37] Do CB. The multivariate Gaussian distribution. Section Notes, Lecture on Machine Learning, CS. 2008;229.

[38] Raschka S, Patterson J, Nolet C. Machine learning in python: Main developments and technology trends in data science, machine learning, and artificial intelligence. Information. 2020;11:193.

[39] Callen HB. Thermodynamics and an introduction to thermostatistics. Second ed: Wiley; 1985.

[40] Vatansever S, Gümüş ZH, Erman B. Intrinsic K-Ras dynamics: A novel molecular dynamics data analysis method shows causality between residue pair motions. Scientific reports. 2016;6:37012.

[41] Zhu K, Li C, Wu KY, Mohr C, Li X, Lanman B. Modeling receptor flexibility in the structure-based design of KRASG12C inhibitors. Journal of Computer-Aided Molecular Design. 2022;36:591–604.

[42] Kapoor A, Travesset A. Differential dynamics of RAS isoforms in GDP-and GTP-bound states. Proteins: Structure, Function, and Bioinformatics. 2015;83:1091–106.

[43] Shen C, Yin J, Wang M, Yu Z, Xu X, Zhou Z, et al. Mutations influence the conformational dynamics of the GDP/KRAS complex. Journal of Biomolecular Structure and Dynamics. 2024:1–14.

[44] Seifert E. OriginPro 9.1: scientific data analysis and graphing software-software review. Journal of chemical information and modeling. 2014;54:1552.

[45] Parikh K, Banna G, Liu SV, Friedlaender A, Desai A, Subbiah V, et al. Drugging KRAS: current perspectives and state-of-art review. Journal of hematology & oncology. 2022;15:152.

[46] Ostrem JM, Peters U, Sos ML, Wells JA, Shokat KM. K-Ras (G12C) inhibitors allosterically control GTP affinity and effector interactions. Nature. 2013;503:548–51.

[47] Ali AA, Dorbath E, Stock G. Allosteric communication mediated by protein contact clusters: A dynamical model. arXiv preprint arXiv:240815110.2024.

[48] Haliloglu T, Hacisuleyman A, Erman B. Prediction of allosteric communication pathways in proteins. Bioinformatics. 2022;38:3590–9.

