## Supplementary for "Dynamic Coupling and Entropy Changes in KRAS G12D Mutation: Insights into Molecular Flexibility, Allostery and Function"

^2^ Department of [Computer Engineering](http://comp.ku.edu.tr/), Koc University Rumelifeneri Yolu, Sariyer 34450, Istanbul Turkey

**Numbering system for GNP used in figures in the manuscript**

Atom number and atom type for GNP

**173	P1**

**174	O1**

**175	O2**

**176	O3**

**177	N1**

**178	P2**

**179	O4**

**180	O5**

**181	O6**

**182	P3**

**183	O7**

**184	O8**

**185	O9**

**186	C1**

**187	C2**

**188	O10**

**189	C3**

**190	O11**

**191	C4**

**192	O12**

**193	C5**

**194	N2**

**195	C6**

**196	N3**

**197	C7**

**198	C8**

**199	O13**

**200	N4**

**201	C9**

**202	N5**

**203	N6**

**204	C10**

**205	H1**

**206	H2**

**207	H3**

**208	H4**

**209	H5**

**210	H6**

**211	H7**

**212	H8**

**213	H9**

**214	H10**

**215	H11**

**216	H12**

**217	H13**

**218	H14**

**Determination of allosteric paths in KRAS**

Overlap figure, Figure S1, of contact map generated by identifying residue pairs within a cutoff distance of 7.3 Å, and of residue pairs with MI values greater than 0.5.

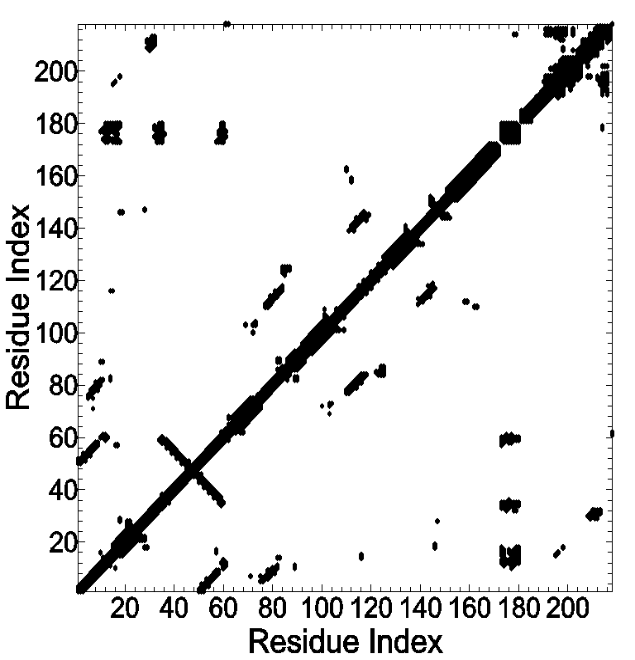

Figure S1. Heat map with cutoff value of 7.3 and MI values larger than 0.5.

**Convergence of trajectories**

Here we show that MI profiles obtained by using 100 and 600 ns trajectories are converged to within 7% of each other, with all peaks in agreement.

In Figure S2, the solid line shows the MI profile obtained by determining KDE using a 600 ns trajectory. The thin line is the profile obtained from a 100 ns trajectory, showing that 100 ns is sufficient for the convergence of mutual information values.

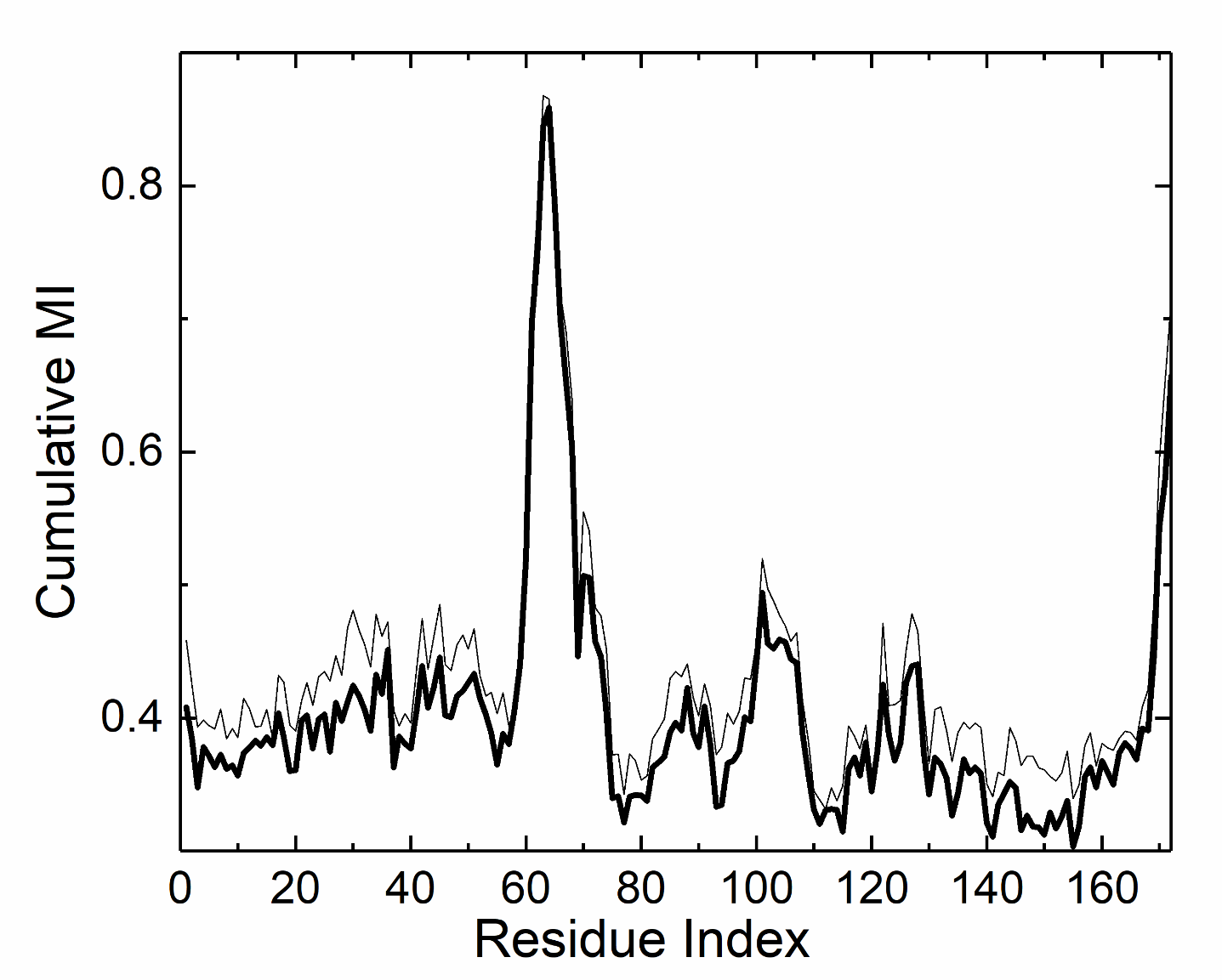

Figure S2. Cumulative mutual information profiles for a 600 ns trajectory (Dark solid line) and for a 100 ns trajectory (thin line).

**Ramachandran Angles for I36 for WT and G12D**

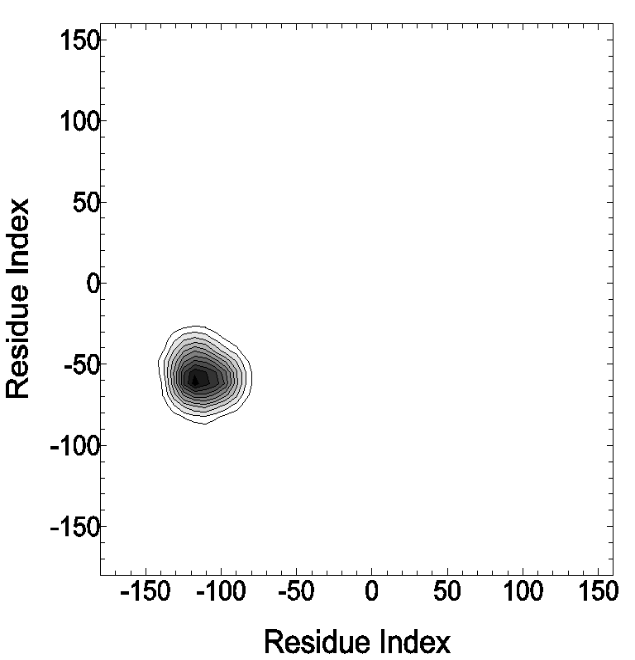

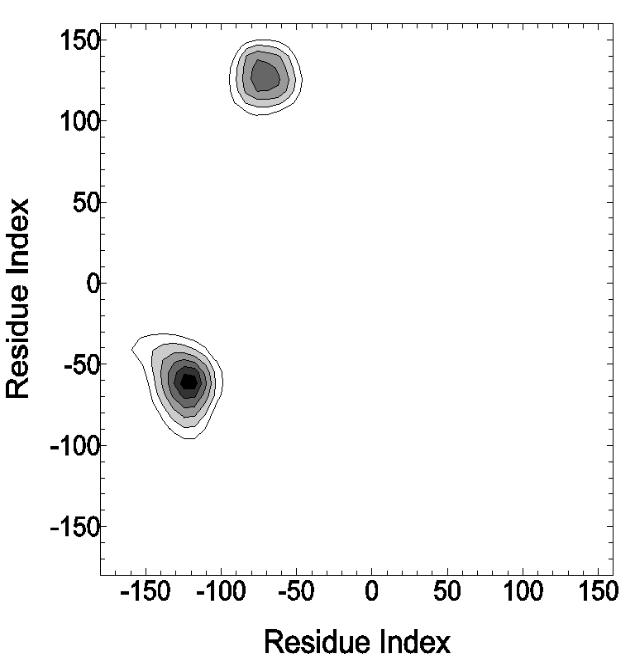

Figure S3. Ramachandran plots of residue I36 from WT (left panel) and G12D (right panel) simulations.

**Ramachandran Angles for G60 for WT and G12D**

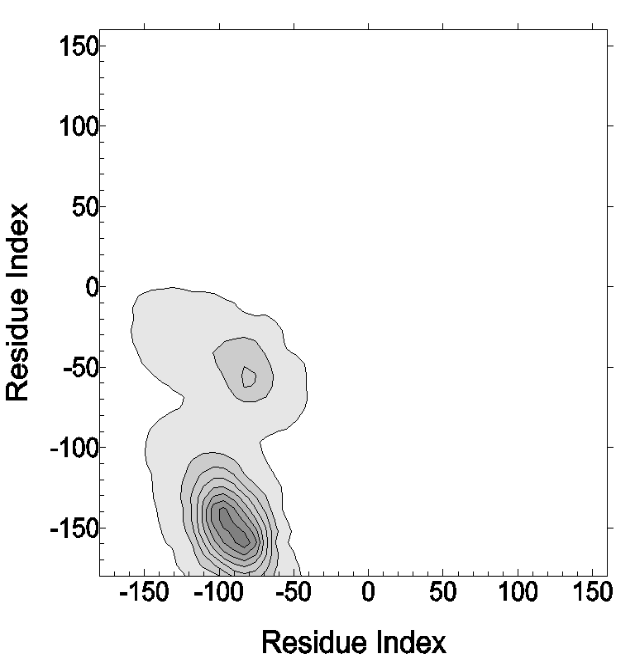

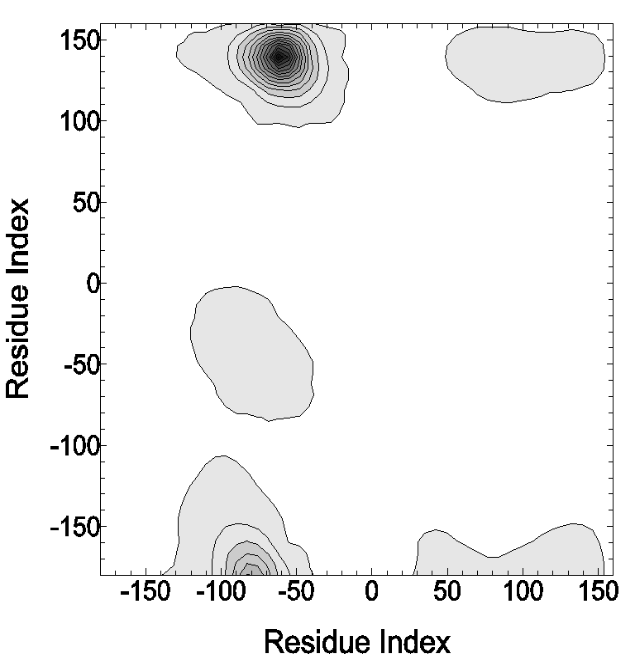

Figure S4. Ramachandran plots of residue G60 from WT (left panel) and G12D (right panel) simulations.

**g) Interactions of GNP with KRAS**

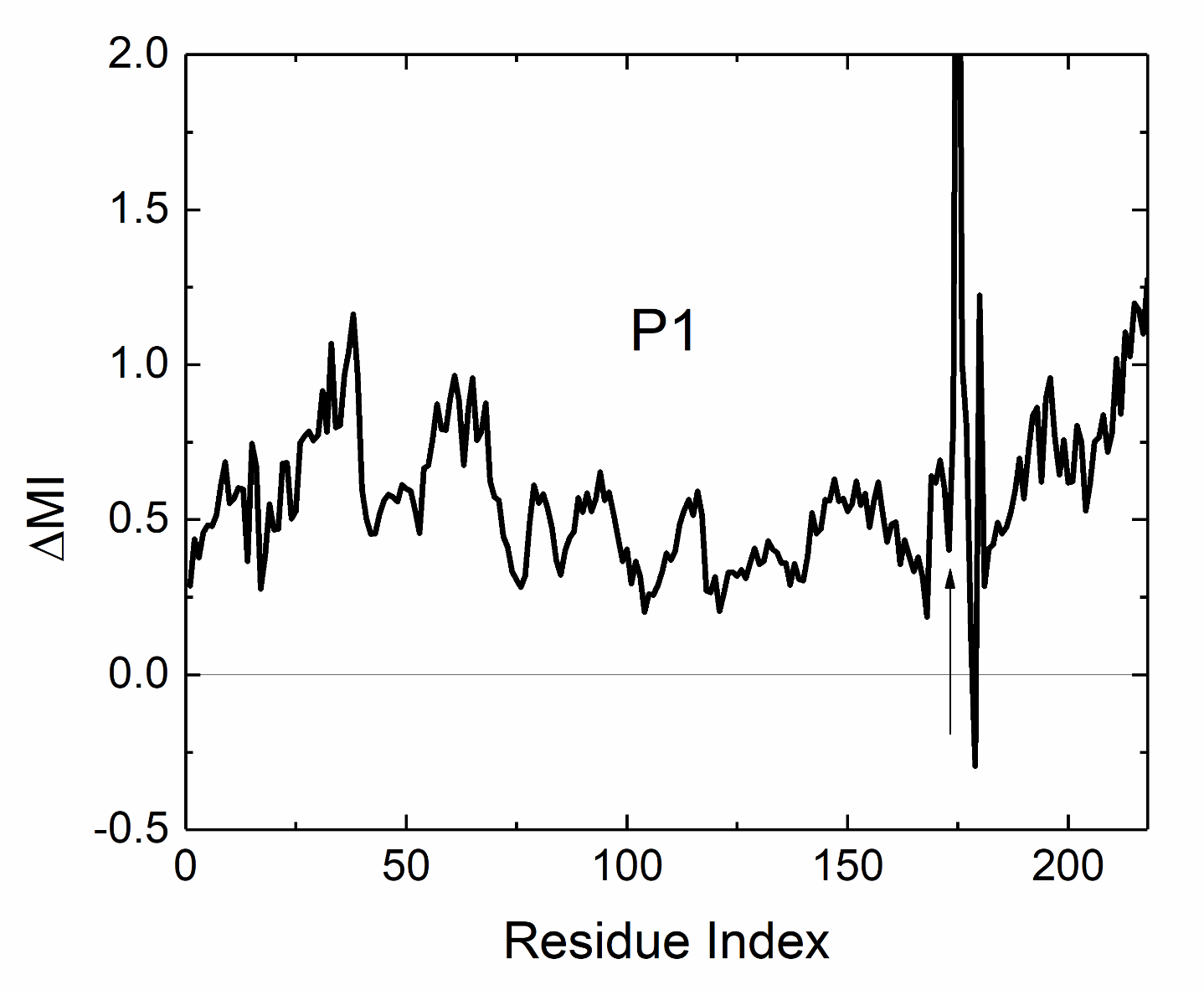

Figure S5. The change of mutual information values of γ-phosphate of GNP with the other residues and GNP atoms. Residues from 1 to 172 belong to KRAS and those larger than 172 refer to the atoms of GNP.

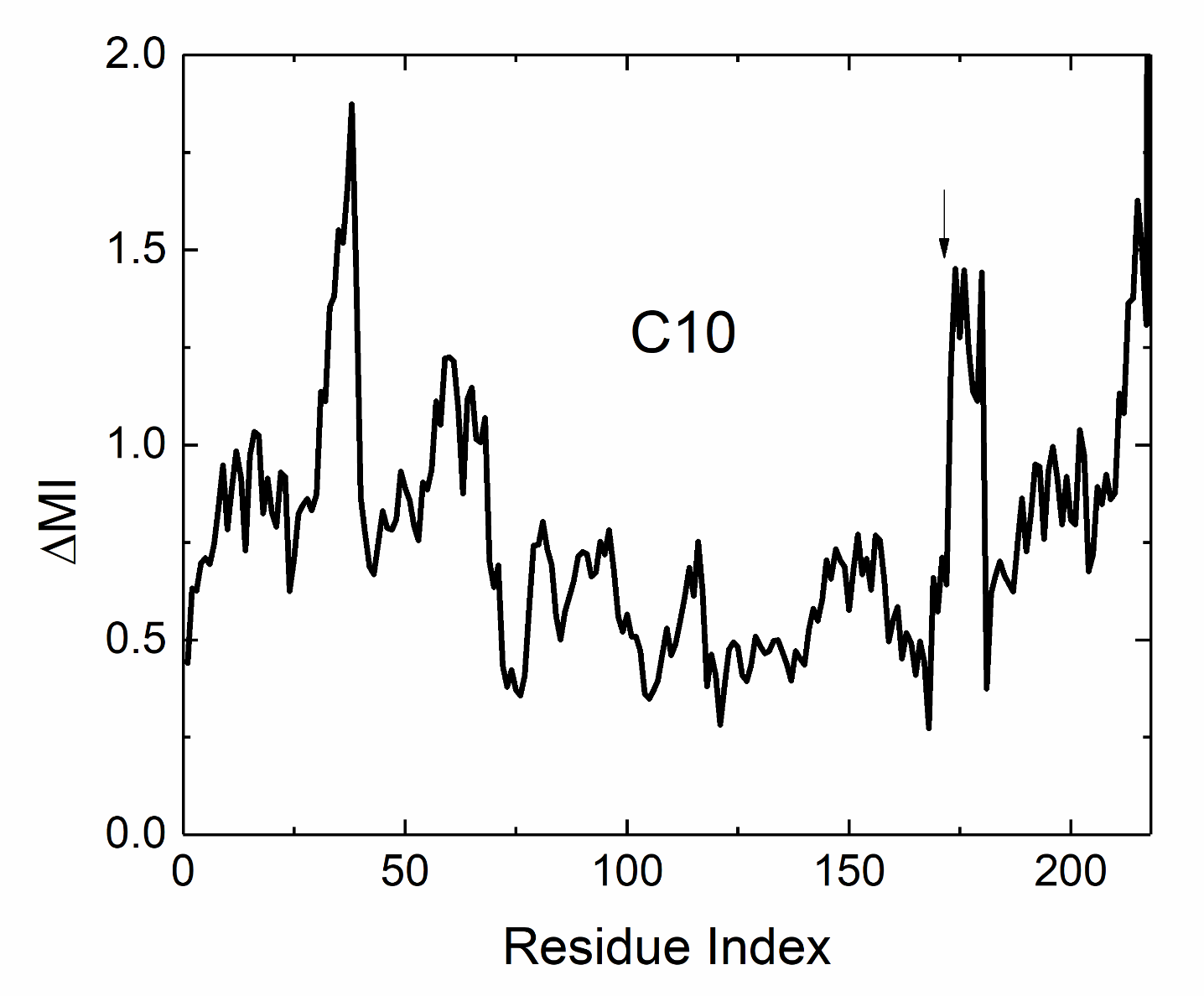

Figure S6. The change of mutual information values of C10 atom of GNP with the other residues and GNP atoms. Residues from 1 to 172 belong to KRAS and those larger than 172 refer to the atoms of GNP.

The condition ΔMI=ΔH(i)+ΔH(j)−ΔH(i,j)>0 or equivalently ΔH(i)+ΔH(j)>ΔH(i,j) reveals an important aspect of how the mutation affects the interactions between residue pairs in KRAS.
